# Scalable, ultra-fast, and low-memory construction of compacted de Bruijn graphs with Cuttlefish 2

**DOI:** 10.1101/2021.12.14.472718

**Authors:** Jamshed Khan, Marek Kokot, Sebastian Deorowicz, Rob Patro

## Abstract

The de Bruijn graph is a key data structure in modern computational genomics, and construction of its compacted variant resides upstream of many genomic analyses. As the quantity of genomic data grows rapidly, this often forms a computational bottleneck.

We present Cuttlefish 2, significantly advancing the state-of-the-art for this problem. On a commodity server, it reduces the graph construction time for 661K bacterial genomes, of size 2.58Tbp, from 4.5 days to 17–23 hours; and it constructs the graph for 1.52Tbp white spruce reads in ∼10 hours, while the closest competitor requires 54–58 hours, using considerably more memory.

## 1. Background

Rapid developments in the throughput and affordability of modern sequencing technologies have made the generation of billions of short-read sequences from a panoply of biological samples highly time- and cost-efficient. The National Center for Biotechnology Information (NCBI) has now moved the Sequence Read Archive (SRA) to the cloud, and this repository stores more than 14 petabytes worth of sequencing data (NCBI Insights). Yet, this is only a fraction of the total sequencing data that has been produced, which is expected to reach exabyte-scale within the current decade (2). In addition to the continued sequencing of an ever-expanding catalog of various types and states of tissues from reference organisms, metagenomic sequencing of environmental (3) and microbiome (4) samples is also expected to enjoy a similar immense growth.

Given the expansive repository of existing sequencing data and the rate of acquisition, Muir et al. (5) argue that the ability of computational approaches to keep pace with data acquisition has become one of the main bottlenecks in contemporary genomics. These needs have spurred methods developers to produce ever more efficient and scalable computational methods for a variety of genomics analysis tasks, from genome and transcriptome assembly to pan-genome analysis. Against this backdrop, the de Bruijn graph, along with its variants, has become a compact and efficient data representation of increasing importance and utility across computational genomics.

The de Bruijn graph originated in combinatorics as a mathematical construct devised to prove a conjecture about binary strings posed by Ir. K. Posthumus (6, 7). In bioinformatics, de Bruijn graphs were introduced in the context of genome assembly algorithms for short-reads (8, 9), although the graph introduced in this context adopts a slightly different definition than in combinatorics. Subsequently, the de Bruijn graph has gradually been used in an increasing variety of different contexts within computational biology, including but not limited to: read correction (10, 11), genomic data compression (12), genotyping (13), structural variant detection (14), read mapping (15, 16), sequence-similarity search (17), metagenomic sequence analysis (18–20), transcriptome assembly (21, 22), transcript quantification (23), and long-read assembly (24–26).

In the context of fragment assembly—whether in forming contigs for whole-genome assembly pipelines (27, 28), or in encapsulating the read set into a summary representative structure for a host of downstream analyses (29–32)—de Bruijn graphs continue to be used extensively. The non-branching paths in de Bruijn graphs are uniquely-assemblable contiguous sequences (known as *unitigs*) from the sequencing reads. Thus, they are certain to be present in any faithful genomic reconstruction from these reads, have no ambiguities regarding repeats in the data, and are fully consistent with the input. As such, maximal unitigs are excellent candidates to summarize the raw reads, capturing their essential substance, and are usually the output of the initial phase of modern *de novo* short-read assembly tools. Collapsing a set of reads into this compact set of fragments that preserve their effective information can directly contribute to the efficiency of many downstream analyses over the read set.

When constructed from reference genome sequences, the unitigs in the de Bruijn graphs correspond to substrings in the references that are shared identically across subsets of the genomes. Decomposing the reference collection into these fragments retains much of its effective information, while typically requiring much less space and memory to store, index, and analyze, than processing the collection of linear genomes directly. The ability to compactly and efficiently represent shared sequences has led many modern sequence analysis tools to adopt the de Bruijn graph as a central representation, including sequence indexers (33), read aligners (15, 16), homology mappers (34, 35), and RNA-seq analysis tools (23, 36, 37). Likewise, pan-genome analysis tools (38–43) frequently make use of the maximal unitigs of the input references as the primary units upon which their core data structures and algorithms are built.

The vast majority of the examples described above make use of the *compacted* de Bruijn graph. A de Bruijn graph is compacted by collapsing each of its maximal, non-branching paths (unitigs) into a single vertex. Many computational genomics workflows employing the (compacted) de Bruijn graph are multi-phased, and typically, their most resource-intensive step is the initial one: construction of the regular and/or the compacted de Bruijn graph. The computational requirements for constructing the graph are often considerably higher than the downstream steps—posing major bottlenecks in many applications (13, 30). As such, there has been a concerted effort over the past several years to develop resource-frugal methods capable of constructing the compacted graph (44–51). Critically, solving this problem efficiently and in a context independent from any specific downstream application yields a modular tool (45, 47) that can be used to enable a wide variety of subsequent computational pipelines.

To address the scalability challenges of constructing the compacted de Bruijn graph, we recently proposed a novel algorithm, Cuttlefish (44), that exhibited faster performance than pre-existing state-of-the-art tools, using (often multiple times) less memory. However, the presented algorithm is only applicable when constructing the graph from existing reference sequences. It cannot be applied in a number of contexts, such as fragment assembly or contig extraction from raw sequencing data. In this paper, we present a fast and memory-frugal algorithm for constructing compacted de Bruijn graphs, Cuttlefish 2, applicable *both* on raw sequencing short-reads and assembled references, that can scale to very large datasets. It builds upon the novel idea of modeling de Bruijn graph vertices as Deterministic Finite Automata (DFA) (52) from Khan and Patro (44). However, the DFA model itself has been modified, and the algorithm has been generalized, so as to accommodate all valid forms of input. At the same time, in the case of constructing the graph from reference sequences, it is considerably faster than the previous approach, while retaining its frugal memory profile. We evaluated Cuttlefish 2 on a collection of datasets with diverse characteristics, and assess its performance compared to other leading compacted de Bruijn graph construction methods. We observed that Cuttlefish 2 demonstrates superior performance in all the experiments we consider.

Additionally, we demonstrate the flexibility of our approach by presenting another application of the algorithm. The compacted de Bruijn graph forms a vertex-decomposition of the graph, while preserving the graph topology (47). However, for some applications, only the vertex-decomposition is sufficient, and preservation of the topology is redundant. For example, for applications such as performing presence-absence queries for k-mers or associating information to the constituent k-mers of the input (53, 54), any set of strings that preserves the exact set of k-mers from the input sequences can be sufficient. Relaxing the defining requirement of unitigs, that the paths be non-branching in the underlying graph, and seeking instead a set of maximal non-overlapping paths covering the de Bruijn graph, results in a more compact representation of the input data. This idea has recently been explored in the literature, with the representation being referred to as a spectrum-preserving string set (55), and the paths themselves as simplitigs (56). We demonstrate that Cuttlefish 2 can seamlessly extract such maximal path covers by simply constraining the algorithm to operate on some specific subgraph(s) of the original graph. We compared it to the existing tools available in the literature (57) for constructing this representation, and observed that it outperforms those in terms of resource requirements.

## 2. Results

### 2.1. Cuttlefish 2 overview

We present a high-level overview of the Cuttlefish 2 algorithm here. A complete treatment is provided in Sec. 3.3.

Cuttlefish 2 takes as input a set ℛ of strings, that are either short-reads or whole-genome references, a k-mer length k, and a frequency threshold f_0_ ≥ 1. As output, it produces the maximal unitigs of the de Bruijn graph G(ℛ, k). Fig. 1 highlights the major steps in the algorithm.

**Figure 1:**
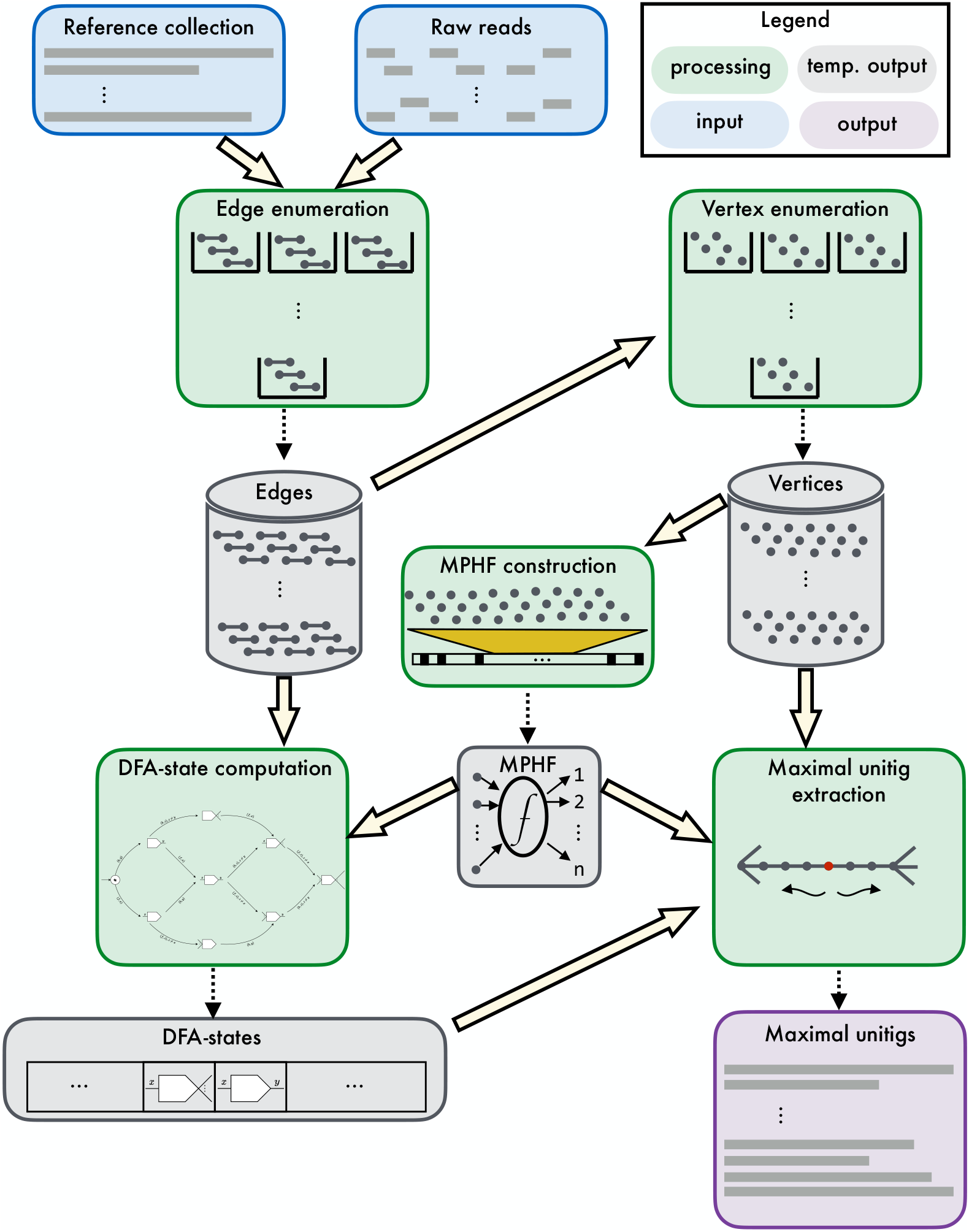
An overview of the Cuttlefish 2 algorithm. It is capable of constructing the compacted de Bruijn graph from a collection of either reference sequences or raw sequencing reads. The edges ((k + 1)-mers) of the underlying de Bruijn graph are enumerated from the input, and optionally filtered based on the user-defined threshold. The edges are then used to enumerate the vertices (k-mers) they contain. An MPHF is constructed over the set of vertices, to associate the DFA-state of each vertex to it. Then the edge set is iterated over to determine the state of the DFA of each vertex in the graph, by transitioning the DFA through appropriate states, based on the edges in which the vertex is observed. Then an iteration over the original vertices to stitch together appropriate edges allows the extraction of the maximal unitigs.

Cuttlefish 2 first enumerates the set ℰ of edges of G(ℛ, k), the (k + 1)-mers present at least f_0_ times in ℛ. This way the potential sequencing errors, present in case in which read sets are given as input, are discarded. Then the set 𝒱 of vertices of G(ℛ, k), which are the k-mers present in these (k + 1)-mers, are extracted from ℰ. Next, a Minimal Perfect Hash Function (MPHF) 𝔣 over these vertices is constructed, that maps them bijectively to [1, |𝒱|]. This provides a space-efficient way to associate information to the vertices through hashing. Modeling each vertex *ν* ∈ 𝒱 as a Deterministic Finite Automaton (DFA), a piecewise traversal on G(ℛ, k) is made using ℰ, computing the state S_*ν*_ of the automaton of each *ν* ∈ 𝒱—associated to *ν* through 𝔣(*ν*). The DFA modeling scheme ensures the retention of just enough information per vertex, such that the maximal unitigs are constructible afterwards from the automata states. Then, with another piece-wise traversal on G(ℛ, k) using 𝒱 and the states collection S, Cuttlefish 2 retrieves all the non-branching edges of G(ℛ, k)—retained by the earlier traversal—and stitches them together in chains, constructing the maximal unitigs.

### 2.2. Experiments

We performed a number of experiments to characterize the various facets of the Cuttlefish 2 algorithm, its implementation, and some potential applications. We evaluated its execution performance compared to other available implementations of leading algorithms on de Bruijn graphs solving—(1) the compacted graph construction and (2) the maximal path cover problems, applicable on shared-memory multi-core machines. Although potentially feasible, Cuttlefish 2 is not designed as a method to leverage the capability of being distributed on a cluster of compute-nodes. Therefore, we did not consider relevant tools operating in that paradigm. We assessed its ability to construct compacted graphs and path covers for both sequencing reads and large pan-genome collections. By working on the (k + 1)-mer spectrum, the new method performs a substantial amount of data reduction on the input sequences, yielding considerable speedups over the Cuttlefish algorithm (44) that, instead, requires multiple passes over the input sequences.

Next, we assess some structural characteristics of the algorithm and its implementation. Given an input dataset and a fixed internal parameter γ, the time- and the space-complexity of Cuttlefish 2 depend on k (see Sec. 3.4). We evaluated the impact of k on its execution performance, and also assessed some structural properties of the compacted graph that change with the parameter k. Moreover, we appraised the parallel scalability of the different steps of the algorithm, characterizing the ones that scale particularly well with increasing processor-thread count, as well as those that saturate more quickly.

A diverse collection of datasets has been used to conduct the experiments. We delineate the pertinent datasets for the experiments in their corresponding sections. The commands used for executing the different tools are available in Suppl. Sec. 1.10.

We compared the outputs of Cuttlefish 2 to those of several other tools used throughout our experiments. A detailed discussion of this is present at Suppl. Sec. 1.5.

#### Computation system for evaluation

All experiments were performed on a single server with two Intel Xeon E5-2699 v4 2.20 GHz CPUs having 44 cores in total and enabling up-to 88 threads, 512 GB of 2.40 GHz DDR4 RAM, and a number of 3.6 TB Toshiba MG03ACA4 ATA HDDs. The system is run with Ubuntu 16.10 GNU/Linux 4.8.0-59-generic. The running times and the maximum memory usages were measured with the GNU `time` command, and the intermediate disk-usages were measured using the Linux commands `inotifywait` and `du`.

### 2.3. Compacted graph construction for sequencing data

We evaluated the performance of Cuttlefish 2 in constructing compacted de Bruijn graphs from short-read sequencing data compared to available implementations of other leading compaction algorithms: (1) ABySS-Bloom-dBG, the maximal unitigs assembler of the ABySS 2.0 assembly-pipeline (27), (2) Bifrost (45), (3) deGSM (46), and (4) BCALM 2 (47).

The performances were tested on a number of short-read datasets with varied characteristics: (1) Mammalian dataset: (i) a human read set (NIST HG004) from an Ashkenazi white female *Homo Sapiens* (paired-end 250 bp Illumina reads with 70x coverage, SRA3440461–95, 148 GB compressed FASTQ), from Zook et al. (58); (ii) an RNA sequencing dataset (ENA PRJEB3365) of 465 human lymphoblastoid cell line samples from the 1000 Genomes project (single-end 36 bp small-RNA-seq Illumina reads, ERP001941, 140 GB compressed FASTQ), from Lappalainen et al. (59). (2) Metagenomic datasets: (i) a gut microbiome read set (ENA PRJEB33098) from nine individuals (paired-end 150 bp Illumina reads with high coverage, ERP115863, 45 GB compressed FASTQ), from Mas-Lloret et al. (60); and (ii) a soil metagenome read set (Iowa Corn) from 100-years-cultivated Iowa agricultural corn soil (paired-end 76 bp and 114 bp Illumina reads with low coverage, SRX100357 and SRX099904–06, 152 GB compressed FASTQ), used by Howe et al. (61); and (3) Large organism dataset: a white spruce read set (NCBI PRJNA83435) from a Canadian *Picea glauca* tree (paired-end 150 bp and 100 bp Illumina reads with high coverage, SRA056234, 1.14 TB compressed FASTQ), from Birol et al. (62). Table 1 contains the summary results of the benchmarking.

**Table 1:**
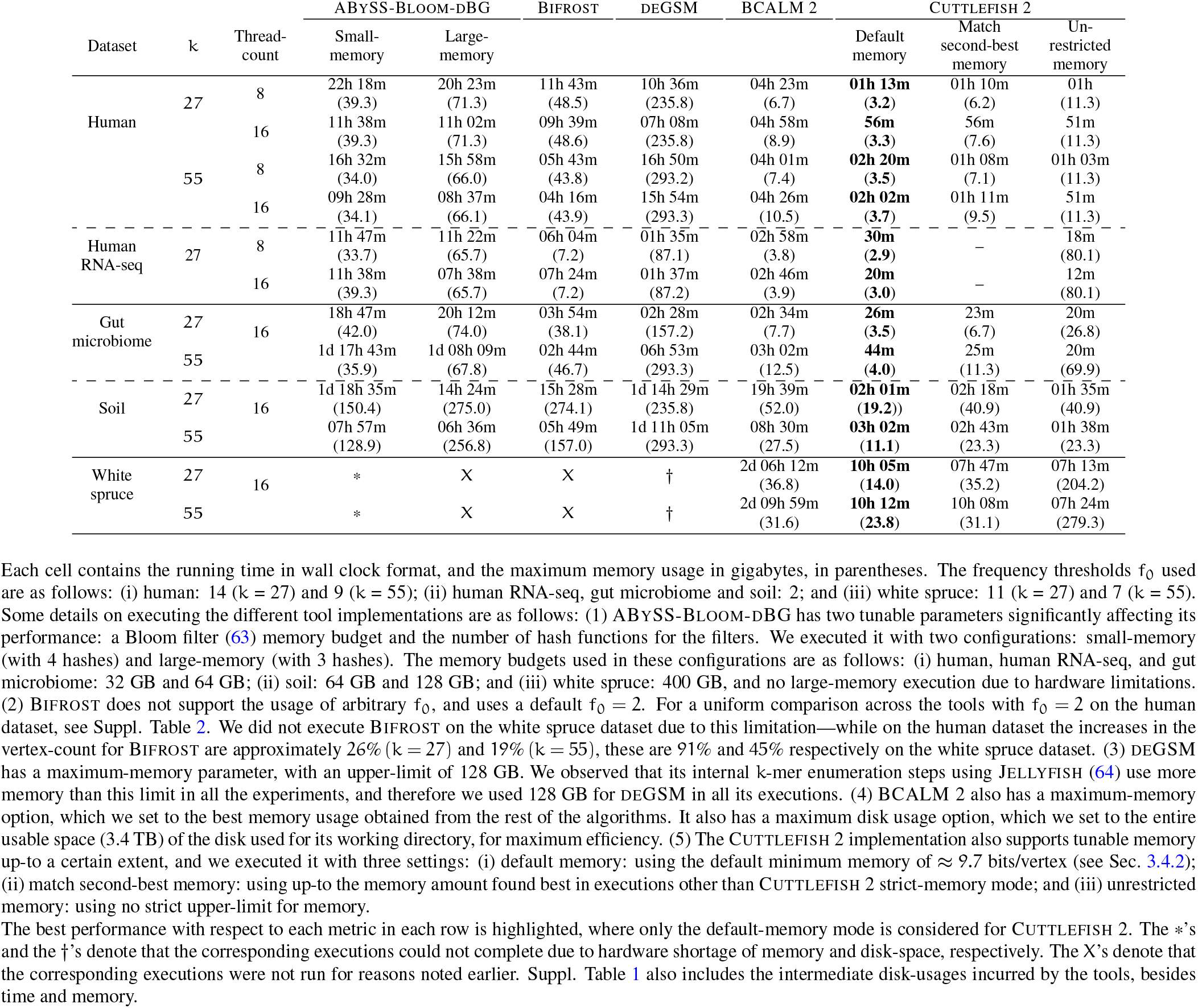
Time- and memory-performance results for constructing compacted de Bruijn graphs from short-read sets.

The frequency threshold f_0_ of k-mers ((k + 1)-mers in case of Cuttlefish 2 ^1^) for the algorithms was approximated using k-mer frequency distributions so as to roughly minimize the misclassification rates of *weak* and *solid* k-mers ^2^ in these experiments (See Suppl. Sec. 1.1). In many practical scenarios, it might be preferable to skip computing an (approximate) frequency distribution, setting f_0_ through some informed choice based on the properties of the input data (e.g. the sequencing depth and protocol). This can incorporate more weak k-mers into the graph. We present the results for such a scenario in Suppl. Table 2 on the human read set, setting f_0_ to just 2.

Across the different datasets and algorithms evaluated, several trends emerge, notable from Table 1. First, we observe that for every dataset considered, Cuttlefish 2 is the fastest tool to process the data, while *simultaneously* using the least amount of memory. If we allow Cuttlefish 2 to match the memory used by the second most memory-frugal method (which is always BCALM 2 here), then it often completes even more quickly. We note that Cuttlefish 2 retains its performance lead over the alternative approaches across a wide range of different data input characteristics.

Among all the methods tested, Cuttlefish 2 and BCALM 2 were the only tools able to process all the datasets to completion under the different configurations tested, within the memory and disk-space constraints of the testing system. The rest of the methods generally required substantially more memory, sometimes over an order of magnitude more, depending on the dataset.

Of particular interest is Cuttlefish 2’s performance compared against BCALM 2. Relative to BCALM 2, Cuttlefish 2 is 1.7–5.3x faster on the human read set, while using 2.1–2.8x less memory. On the RNA-seq dataset, it is 8.3– 5.9x faster, with 1.3x less memory. For the metagenomic datasets, it is 4.1–5.9x faster and uses 2.2–3.1x less memory on the gut microbiome data, and is 2.8–8.5x faster using 2.5– 2.7x less memory on the soil data. On the largest sequencing dataset here, the white spruce read set, Cuttlefish 2 is 5.4– 5.7x faster and is 1.3–2.6x memory-frugal—taking about 10 hours, compared to at least 54 hours for BCALM 2.

The timing-profile of BCALM 2 and Cuttlefish 2 excluding their similar initial stage: k-mer and (k + 1)-mer enumeration, respectively, are shown in Suppl. Table 4. We also note some statistics of the de Bruijn graphs and their compacted forms for these datasets in Suppl. Table 5.

### 2.4. Compacted graph construction for reference collections

We assessed the execution performance of Cuttlefish 2 in constructing compacted de Bruijn graphs from whole-genome sequence collections in comparison to the available implementations of the following leading algorithms: (1) Bifrost (45), (2) deGSM (46), and (3) BCALM 2 (47). TwoPaCo (48) is another notable algorithm applicable in this scenario, but we did not include it in the benchmarking as its output step lacks a parallelized implementation, and we consider very large sequence collections in this experiment.

We evaluated the performances on a number of datasets with varying attributes: (1) Metagenomic collection: 30,691 representative sequences from the most prevalent human gut prokaryotic genomes, gathered by Hiseni et al. (66) (≈ 61B bp, 18 GB compressed FASTA); (2) Mammalian collection: 100 human genomes—7 real sequences from Baier et al. (49) and 93 sequences simulated by Minkin et al. (48) (≈ 294B bp, 305 GB uncompressed FASTA); and (3) Bacterial archive: 661,405 bacterial genomes, collected by Blackwell et al. (67) from the European Nucleotide Archive (≈ 2.58T bp, 752 GB compressed FASTA). Table 2 conveys the summary results of the benchmarking.

**Table 2:**
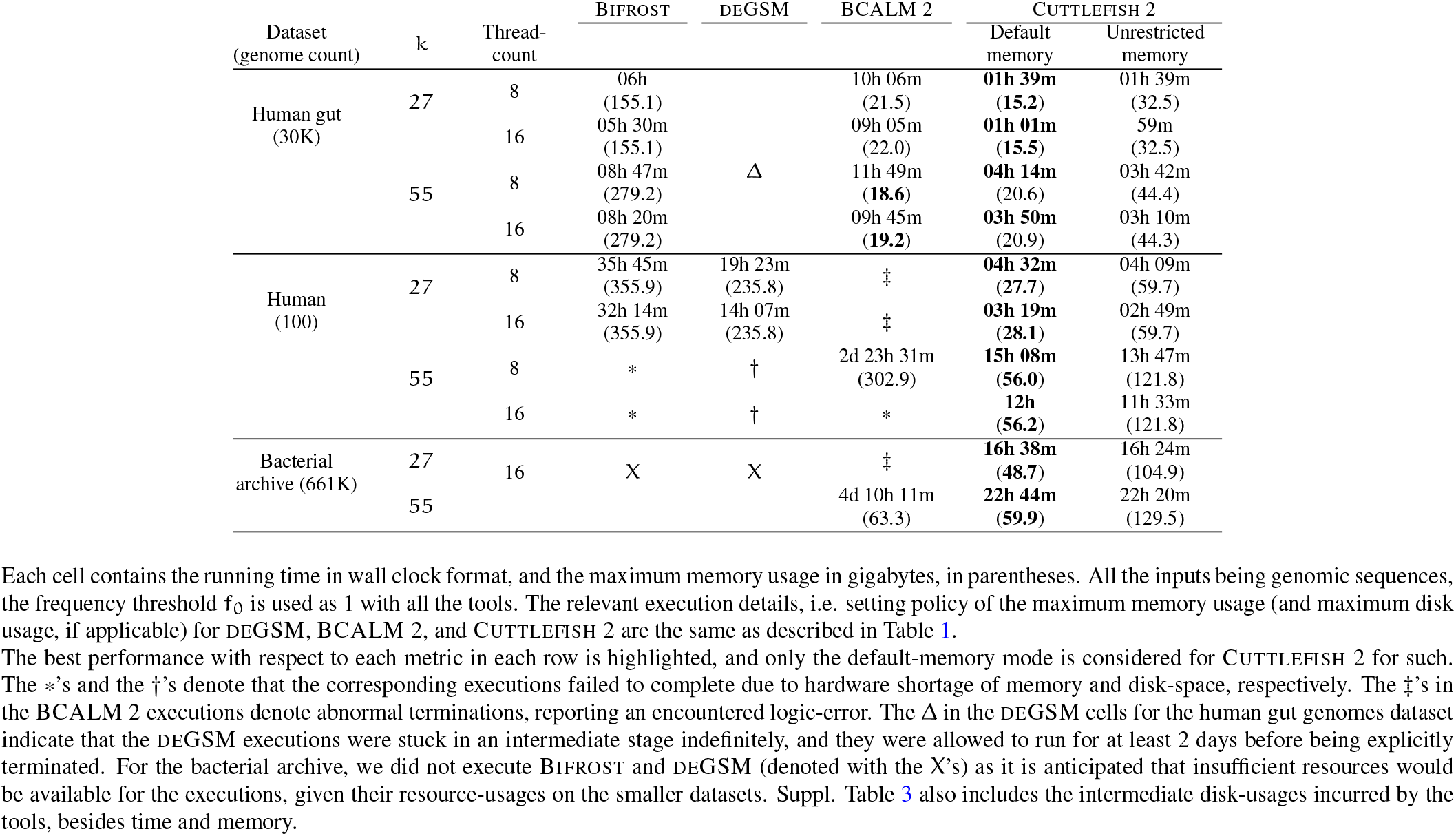
Time- and memory-performance results for constructing compacted de Bruijn graphs from whole-genome reference collections.

Evaluating the performance of the different tools over these pan-genomic datasets, we observe similar trends to what was observed in Table 1, but with even more extreme differences than before. For a majority of the experiment configurations here, only BCALM 2 and Cuttlefish 2 were able to finish processing within time- and machine-constraints. Again, Cuttlefish 2 exhibits the fastest runtime on all datasets, and the lowest memory usage on all datasets except the human gut genomes (where it consumes 1–2 GB more memory than BCALM 2, though taking 6–7 fewer hours to complete).

Cuttlefish 2 is 2.4–8.9x faster on the 30K human gut genomes compared to the closest competitors, using similar memory. On the 100 human reference sequences, Cuttlefish 2 is 4.3–4.7x faster, using 5.4–8.4x less memory. The only other tools able to construct this compacted graph successfully are deGSM for k = 27 (taking 4.3x as long and requiring 8.4x as much memory as Cuttlefish 2) and BCALM 2 for k = 55 (taking over 4.7x as long and 5.4x as much memory as Cuttlefish 2). Finally, when constructing the compacted graph on the 661,405 bacterial genomes, Cuttlefish 2 is the only tested tool able to construct the graph for k = 27. For k = 55, BCALM 2 also completed, taking about 4.5 days, while Cuttlefish 2 finished under a day, with similar memory-profile. Overall, we observe that for large pan-genome datasets, Cuttlefish 2 is not only considerably faster and more memory-frugal than alternative approaches, but is the only tool able to reliably construct the compacted de Bruijn graph under all the different configurations tested, within the constraints of the experimental system.

Table 4 notes the timing-profiles for BCALM 2 and Cuttlefish 2 without their first step of k-mer and (k + 1)-mer enumerations, and Table 5 shows some characteristics of the (compacted) de Bruijn graphs for these pan-genome datasets.

### 2.5. Maximal path cover construction

The execution performance of Cuttlefish 2 in decomposing de Bruijn graphs into maximal vertex-disjoint paths was assessed compared to the only two available tool implementations in literature (57) for this task: (1) ProphAsm (56) and (2) UST (55).

For sequencing data, we used: (1) a roundworm read set (ENA DRR008444) from a *Caenorhabditis elegans* nematode (paired-end 300 bp Illumina reads, 5.6 GB compressed FASTQ); (2) the gut microbiome read set (ENA PRJEB33098) noted earlier; and (3) the human read set (NIST HG004) noted earlier. For whole-genome data, we used sequences from: (1) a roundworm reference (*Caenorhabditis elegans*, VC2010) (68); (2) a human reference (*Homo sapiens*, GRCh38); and (3) 7 real humans, collected from Baier et al. (49). Table 3 presents the summary results of the bench-marking.

**Table 3:**
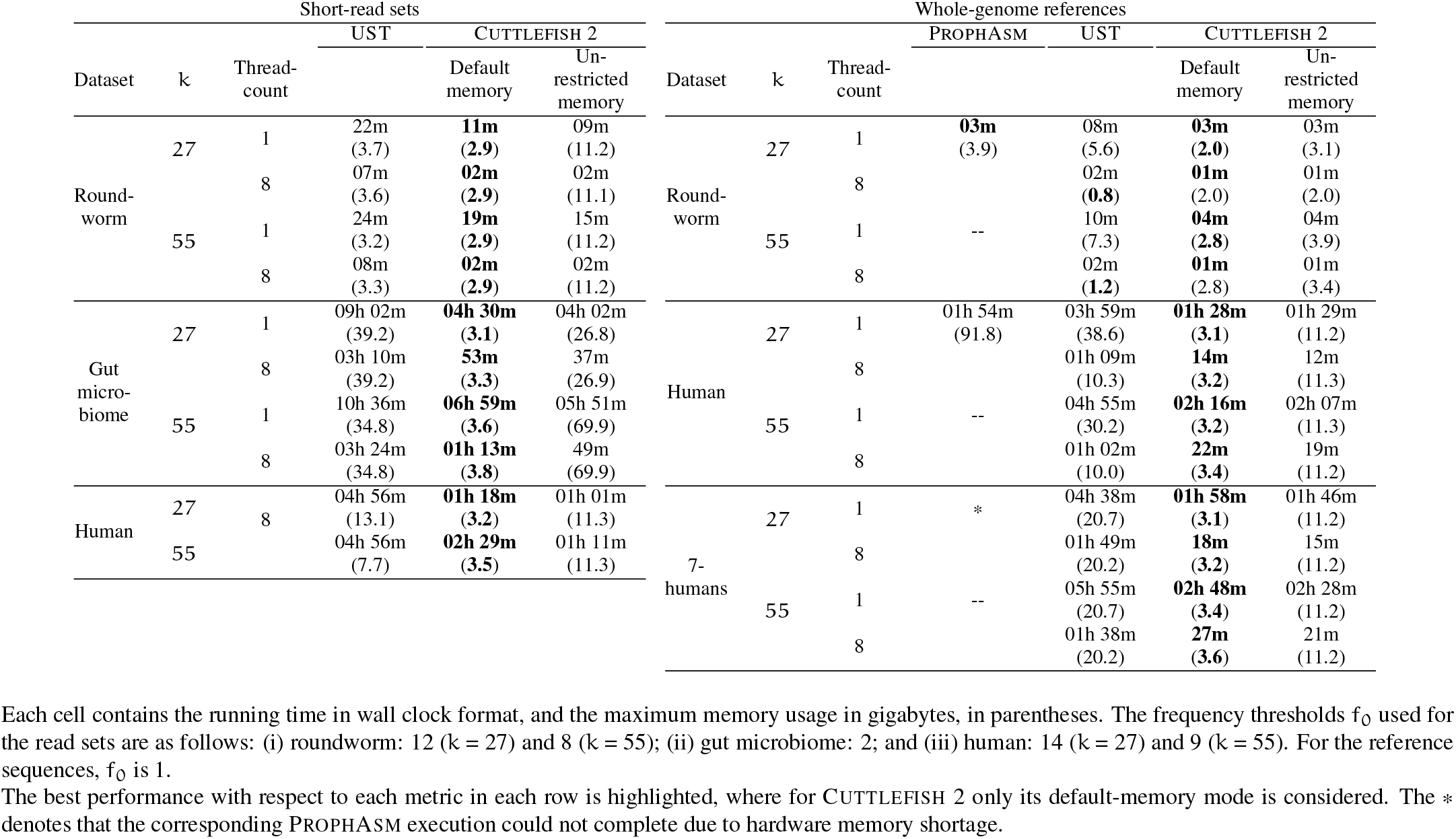
Time- and memory-performance results for decomposing de Bruijn graphs into maximal vertex-disjoint paths.

**Table 4:** Time- and memory-performance of Cuttlefish 2 for constructing the compacted de Bruijn graph from the human read set NIST HG004, and some corresponding metrics of the output maximal unitigs, over a range of k-mer sizes.

| k | k-mer count | Performance-metrics |  | Unitig-metrics |  |  |  |  |
| --- | --- | --- | --- | --- | --- | --- | --- | --- |
|  |  | Default memory | Unrestricted memory | Count | Avg. length (bp) | Max. length (bp) | N50 (bp) | NGA50 (bp) |
| 27 | 2,547,479,119 | 1h 12m (3.19) | 54m (11.29) | 80,465,421 | 58 | 20,648 | 62 | 425 |
| 41 | 2,771,918,177 | 2h 19m (3.48) | 1h 05m (11.26) | 44,768,246 | 102 | 29,381 | 186 | 769 |
| 55 | 2,900,387,834 | 2h 12m (3.54) | 1h 04m (11.28) | 28,510,532 | 156 | 32,725 | 386 | 1,030 |
| 69 | 2,978,629,926 | 2h 42m (3.66) | 1h 11m (19.49) | 20,361,009 | 214 | 45,495 | 552 | 1,256 |
| 83 | 3,029,739,673 | 2h 39m (3.68) | 1h 04m (22.34) | 16,220,627 | 269 | 45,359 | 645 | 1,435 |
| 97 | 3,066,350,056 | 3h 05m (3.78) | 1h 06m (30.57) | 13,938,567 | 316 | 57,338 | 675 | 1,543 |
| 111 | 3,093,353,953 | 2h 53m (3.75) | 1h 08m (32.18) | 12,683,849 | 354 | 57,402 | 660 | 1,596 |
| 125 | 3,111,450,986 | 3h 01m (3.80) | 1h 16m (42.18) | 11,855,026 | 386 | 57,416 | 634 | 1,617 |

We note that Cuttlefish 2 outperforms the alternative tools for constructing maximal path covers in terms of the time and memory required. In the context of this task, Cuttlefish 2 also offers several qualitative benefits over these tools. For example, ProphAsm exposes only a single-threaded implementation. Further, it is restricted to values of k ≤ 32 and only accepts genomic sequences as input (and thus is not applicable for read sets). UST first makes use of BCALM 2 for maximal unitigs extraction—which we observed to be outperformed by Cuttlefish 2 in the earlier experiments— and then employs a sequential graph traversal on the compacted graph to extract a maximal path cover. For this problem, Cuttlefish 2 bypasses the compacted graph construction, and directly extracts a maximal cover.

We observe that compared to the tools, Cuttlefish 2 is competitive on single-threaded executions. While on moderate-sized datasets using multiple threads, it was 2–3.8x faster than UST using 2.2–12.6x less memory on sequencing data, and for reference sequences it was 2.8–6.1x faster than UST using 2.9–6.3x less memory.

We also provide a comparison of the maximal unitig-based and the maximal path cover-based representations of de Bruijn graphs in Suppl. Table 6. We observe that, for the human read set, the path cover representation requires 19– 24% less space than the unitigs. For the human genome reference and 7 humans pan-genome references, these reductions are 14–22%, and 20–25%, respectively. From the statistics of both the representations on the gut microbiome read set, it is evident that the corresponding de Bruijn graphs are highly branching, as might be expected for metagenomic data. The space reductions with path cover in these graphs are 33–36%.

### 2.6. Structural characteristics

Given an input dataset ℛ and a fixed frequency threshold f_0_ for the edges (i.e. (k + 1)-mers), the structure of the de Bruijn graph G(ℛ, k) is completely determined by the k-mer-size—the edge- and the vertex-counts depend on k, and the asymptotic characteristics of the algorithm are dictated only by the k-mer size k and the hash function space-time tradeoff factor γ (see Sec. 3.4). We evaluated how Cuttlefish 2 is affected practically by changes in the k-value. The human read set (NIST HG004) noted earlier was used for these evaluations.

For a range of increasing k-values (and a constant γ), we measured the execution performance of Cuttlefish 2, and the following metrics of the maximal unitigs it produced: the number of unitigs, the average and the maximum unitig lengths, along with the N50 ^3^ and the NGA50 ^4^ scores for contig-contiguity. Across the varying k’s, Table 4 reports the performance- and the unitig-metrics.

The unitig-metrics on this data convey what one might expect—as k increases, so do the average and the maximum lengths of the maximal unitigs, and the N50 and NGA50 metrics, since the underlying de Bruijn graph typically gets less tangled as the k-mer size exceeds repeat lengths (70). It is worth noting that, since we consider here just the extraction of unitigs, and no graph cleaning or filtering steps (e.g. bubble popping and tip clipping), we expect the N50 to be fairly short.

Perhaps the more interesting observation to be gleaned from the results is the scaling behavior of Cuttlefish 2 in terms of k. While the smallest k-value leads to the fastest overall graph construction, with increase in the machine-word count to encode the k-mers, the increase in runtime is rather moderate with respect to k, which follows the expected asymptotics (see Sec. 3.4.1). On the other hand, we observe that this increase can be offset by allowing Cuttlefish 2 to execute with more memory (which helps in the bottleneck step, (k + 1)-mer enumeration). We also note that, while the timing-profile exhibits reasonably good scalability over the parameter k, the effect on the required memory is rather small—it is not directly determined by the k-value, rather is completely dictated by the distinct k-mer count (see Sec. 3.4.2).

### 2.7. Parallel scaling

We assessed the scalability of Cuttlefish 2 across a varying number of processor-threads. For this experiment, we downsampled the human read set NIST HG004 from 70x to 20x coverage and used this as input. We set k to 27 and 55, and executed Cuttlefish 2 with thread-counts ranging in 1–32. For k = 27, Fig. 2a shows the time incurred by each step of the algorithm, and Fig. 2b demonstrates their speedups (i.e. factor of improvement in the speed of execution with different number of processor-threads). Suppl. Fig. 2 shows these metrics for k = 55.

**Figure 2:**
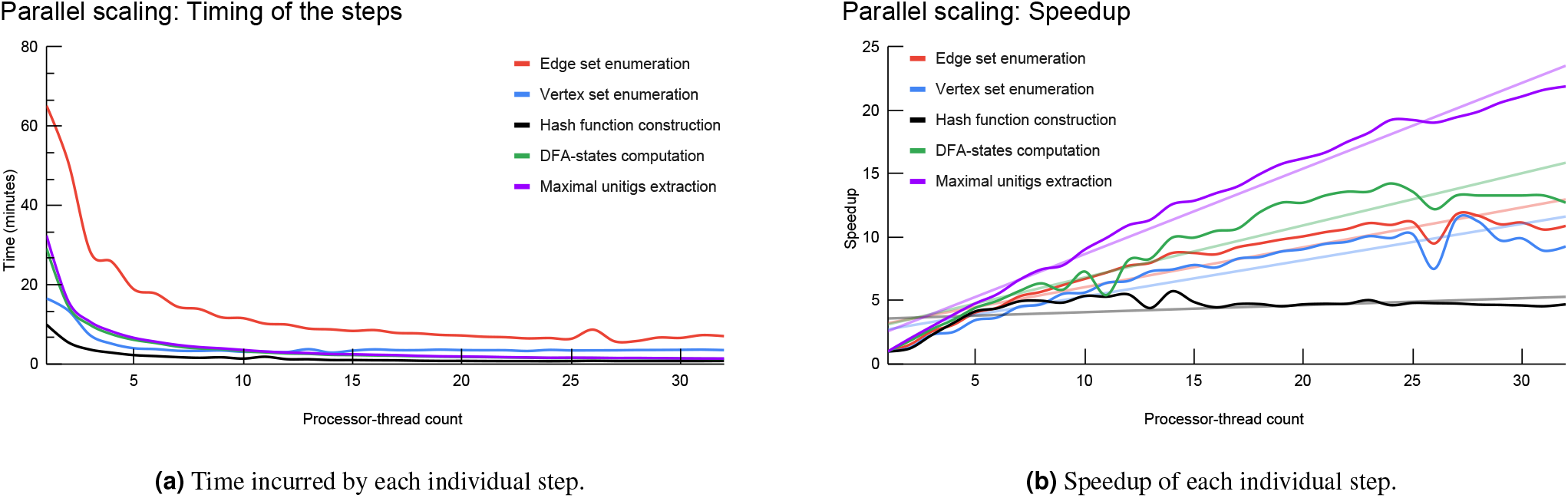
Parallel-scaling metrics for Cuttlefish 2 across 1–32 processor threads, using k = 27 on the (downsampled) human read set NIST HG004, with the frequency threshold f_0_ = 4.

On the computation system used, we observe that all steps of Cuttlefish 2 scale well until about 8 threads. Beyond 8 threads, most steps but the minimal perfect hash construction continue to scale. Fig. 2a shows that the most time-intensive step in the algorithm is the initial edge set enumeration. This step, along with vertex enumeration and DFA states computation, continue to show reasonably good scaling behavior until about 20 threads, then gradually saturating. The final step of unitigs extraction seems to scale well up to the maximum thread-count we tested with (32 in this case).

It is worth reiterating that all experiments were performed on standard hard drives, and that the most resource-intensive step of edge enumeration can be quite input-output (IO) bound, while the rest of the steps also iterate through the indisk set of edges or vertices—bound by disk-read speed. So one might expect different (and quite possibly better) scaling behavior for the IO-heavy operations when executing on faster external storage, e.g. in the form of SSD or NVMe drives (71). This is further evidenced by Kokot et al. (72), who show that KMC 3, the method used for the edge and the vertex enumeration steps in Cuttlefish 2, could have considerable gains in speed on large datasets when executed on SATA SSDs.

In this paper, we present Cuttlefish 2, a new algorithm for constructing the compacted de Bruijn graph, which is very fast and memory-frugal, and highly-scalable in terms of the extent of the input data it can handle. Cuttlefish 2 builds upon the work of Khan and Patro (44), which already advanced the state-of-the-art in reference-based compacted de Bruijn graph construction. Cuttlefish 2 simultaneously addresses the limitation and the bottleneck of Cuttlefish, by substantially generalizing the work to allow graph construction from both raw sequencing reads and reference genome sequences, while offering a more efficient performance profile. It achieves this, in large part, through bypassing the need to make multiple passes over the original input for very large datasets.

As a result, Cuttlefish 2 is able to construct compacted de Bruijn graphs much more quickly, while using less memory—both often multiple *times*—than the numerous other methods evaluated. Since the construction of the graph resides upstream of many computational genomics analysis pipelines, and as it is typically one of the most resource-intensive steps in these approaches, Cuttlefish 2 could help remove computational barriers to using the de Bruijn graph in analyzing the ever-larger corpora of genomic data.

In addition to the advances it represents in the compacted graph construction, we also demonstrate the ability of the algorithm to compute another spectrum-preserving string set of the input sequences—maximal path covers that have recently been adopted in a growing variety of applications in the literature (57). A simple restriction on the considered graph structure allows Cuttlefish 2 to build this construct much more efficiently than the existing methods.

Though a thorough exploration of the potential benefits of improved compacted de Bruijn graph construction to the manifold downstream analyses is outside the scope of the current work, we present a proof of concept application (Suppl. Sec. 1.9), demonstrating the benefits of our improved algorithm to the task of constructing an associative k-mer index. As the scale of the data on which the de Bruijn graph and its variants must be constructed increases, and as the de Bruijn graph itself continues to find ever-more widespread uses in genomics, we anticipate that Cuttlefish 2 will enable its use in manifold downstream applications that may not have been possible earlier due to computational challenges, paving the way for an even more widespread role for the de Bruijn graph in high-throughput computational genomics.

Cuttlefish 2 is implemented in `C++14`, and is available open-source at https://github.com/COMBINE-lab/cuttlefish.

## 3. Methods

### 3.1. Related work

Here we briefly discuss the other compacted de Bruijn graph construction algorithms included in the experiments against which we compare Cuttlefish 2. The BCALM algorithm (50) partitions the k-mers from the input that pass frequency filtering into a collection of disk-buckets according to their minimizers (73), and processes each bucket sequentially as per the minimizer-ordering— loading all the strings of the bucket into memory, joining (or, *compacting*) them maximally while keeping the resulting paths non-branching in the underlying de Bruijn graph, and distributing each resultant string into some other yet-to-be-processed bucket for potential further compaction, or to the final output. As is, BCALM is inherently sequential. BCALM 2 (47) builds upon this use of minimizers to partition the graph, but it modifies the k-mer partitioning strategy so that multiple disk-buckets can be compacted correctly in parallel, and then glues the further compactable strings from the compacted buckets.

ABySS-Bloom-dBG is the maximal unitigs assembler of the ABySS 2.0 assembly tool (27). It first saves all the k-mers from the input reads into a cascading Bloom filter (63) to discard the likely-erroneous k-mers. Then it identifies the reads that consist entirely of retained k-mers, and extends them in both directions within the de Bruijn graph through identifying neighbors using the Bloom filter, while discarding the potentially false-positive paths based on their spans— producing the maximal unitigs.

deGSM first enumerates all the (k + 2)-mers of the input that pass frequency filtering. Then using a parallel external sorting over partitions of this set, it groups together the (k + 2)-mers with the same middle k-mer, enabling it to identify the branching vertices in the de Bruijn graph. Then it merges the k-mers from the sorted buckets in a strategy so as to produce a Burrows-Wheeler Transform (74) of the maximal unitigs.

Bifrost (45) constructs an approximate compacted de Bruijn graph first by saving the k-mers from the input in a Bloom filter (63), and then for each potential non-erroneous k-mer, it extracts the maximal unitig containing it by extending the k-mer in both directions using the Bloom filter. Then using a k-mer counting based strategy, it refines the graph by removing the false edges induced by the Bloom filter.

### 3.2. Definitions

A *string* s is an ordered sequence of symbols drawn from an alphabet Σ. For the purposes of this paper, we assume all strings to be over the alphabet Σ = {A, C, G, T}, the DNA alphabet where each symbol has a reciprocal complement—the complementary pairs being {A, T} and {C, G}. For a symbol c ∈ Σ, 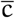 denotes its complement. |s| denotes the length of s. A k*-mer* is a string with length k. s_i_ denotes the i’th symbol in s (with 1-based indexing). A *substring* of s is a string entirely contained in s, and s_i‥j_ denotes the substring of s located from its i’th to the j’th indices, inclusive. pre_𝓁_(s) and suf_𝓁_(s) denote the prefix and the suffix of s with length 𝓁 respectively, i.e. pre_𝓁_(s) = s_1‥𝓁_ and suf_𝓁_(s) = s_|s|−𝓁+1‥|s|_, for some 0 < 𝓁 ≤ |s|. For two strings x and y with suf_𝓁_(x) = pre_𝓁_(y), the 𝓁*-length glue* operation ⊙^𝓁^ is defined as x ⊙^𝓁^ y = x · y_𝓁+1‥|y|_, where denotes the *append* operation.

For a string s, its *reverse complement* 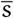 is the string obtained through reversing the order of the symbols in s, and replacing each symbol with its complement, i.e. 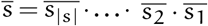. The *canonical form* 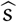 of s is defined as the string 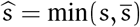, according to some consistent ordering of the strings in Σ^|s|^. In this paper, we adopt the lexicographic ordering of the strings.

Given a set 𝒮 of strings and an integer k > 0, let 𝒦 and 𝒦_+1_ be respectively the sets of k-mers and (k + 1)-mers present as substrings in some s ∈ 𝒮. The (directed) *node-centric de Bruijn graph* G_1_(𝒮, k) = (𝒱_1_, ℰ_1_) is a directed graph where the vertex set is 𝒱_1_ = 𝒦, and the edge set ℰ_1_ is induced by 𝒱_1_: a directed edge e = (u, *ν*) ∈ ℰ_1_ iff suf_k−1_(u) = pre_k−1_(*ν*). The (directed) *edge-centric de Bruijn graph* G_2_(𝒮, k) = (𝒱_2_, ℰ_2_) is a directed graph where the edge set is ℰ_2_ = 𝒦_+1_: each e ∈ 𝒦_+1_ constitutes a directed edge (*ν*_1_, *ν*_2_) where *ν*_1_ = pre_k_(e) and *ν*_2_ = suf_k_(e), and the vertex set 𝒱_2_ is thus induced by ℰ_2_. ^5^

In this work, we adopt the edge-centric definition of de Bruijn graphs. Hence, we use the terms k-mer and vertex and the terms (k + 1)-mer and edge interchangeably. We introduce both variants of the graph here as we compare (in Sec. 2) our algorithm with some other methods that employ the node-centric definition.

We use the *bidirected* variant of de Bruijn graphs in the proposed algorithm, and redefine 𝒦_+1_ to be the set of canonical (k + 1)-mers 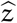 such that z or 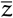 appears as substring in some s ∈ 𝒮. ^6^ For a bidirected edge-centric de Bruijn graph G(𝒮, k) = (𝒱, ℰ) — (i) the vertex set 𝒱 is the set of canonical forms of the k-mers present as substrings in some e ∈ 𝒦_+1_, and (ii) the edge set is ℰ = 𝒦_+1_, where an e ∈ ℰ connects the vertices 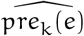 and 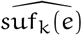. A vertex *ν* has exactly two *sides*, referred to as the *front* side and the *back* side.

For a (k + 1)-mer z such that 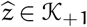, let 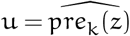 and 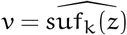 induces an edge from the vertex u to the vertex *ν*, and it is said to *exit* u and *enter ν*. z connects to u’s back iff pre_k_(z) is in its canonical form, i.e. pre_k_(z) = u, and otherwise it connects to u’s front. Conversely, z connects to *ν*’s front iff suf_k_(z) = *ν*, and otherwise to *ν*’s back. Concisely put, z exits through u’s back iff z’s prefix k-mer is canonical, and enters through *ν*’s front iff z’s suffix k-mer is canonical. The edge corresponding to z can be expressed as ((u, ψ_u_),(*ν*, ψ_*ν*_)) : it connects from the side ψ_u_ of the vertex u to the side ψ_*ν*_ of the vertex *ν*.

We prove in Lemma 1 (see Suppl. Sec. 3) that the two (k + 1)-mers z and 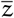 correspond to the same edge, but in reversed directions: 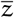 induces the edge ((*ν*, ψ_*ν*_),(u, ψ_u_)) — opposite from that of z. The two edges for z and 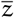 constitute a bidirected edge e = {(u, ψ_u_),(*ν*, ψ_*ν*_)}, corresponding to 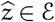. While crossing e during a graph traversal, we say that e *spells* the (k + 1)-mer z when the traversal is from (u, ψ_u_) to (*ν*, ψ_*ν*_), and it spells 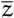 in the opposite direction.

A *walk* w = (*ν*_0_, e_1_, *ν*_1_, …, *ν*_n−1_, e_n_, *ν*_n_) is an alternating sequence of vertices and edges in G(𝒮, k), where the vertices *ν*_i_ and *ν*_i+1_ are connected with the edge e_i+1_, and e_i_ and e_i+1_ are incident to different sides of *ν*_i_. |w| denotes its length in vertices, i.e. |w| = n + 1. *ν*_0_ and *ν*_n_ are its *endpoints*, and the *ν*_i_ for 0 < i < n are its *internal vertices*. The walks (*ν*_0_, e_1_, …, e_n_, *ν*_n_) and (*ν*_n_, e_n_, …, e_1_, *ν*_0_) denote the same walk but in opposite *orientations*. If the edge e_i_ spells the (k + 1)-mer l_i_, then w spells l_1_ ⊙^k^ l_2_ ⊙^k^ … ⊙^k^ l_n_. If |w| = 1, then it spells *ν*_0_. A *path* is a walk without any repeated vertex.

A *unitig* is a path p = (*ν*_0_, e_1_, *ν*_1_, …, e_n_, *ν*_n_) such that either |p| = 1, or in G(𝒮, k):

1. each internal vertex *ν*_i_ has exactly one incident edge at each of its sides, the edges being e_i_ and e_i+1_
2. and *ν*_0_ has only e_1_ and *ν*_n_ has only e_n_ incident to their sides to which e_1_ and e_n_ are incident to, respectively.

A *maximal unitig* is a unitig p = (*ν*_0_, e_1_, *ν*_1_, …, e_n_, *ν*_n_) such that it cannot be extended anymore while retaining itself a unitig: there exists no x, y, e_0_, or e_n+1_ such that (x, e_0_, *ν*_0_, …, e_n_, *ν*_n_) or (*ν*_0_, e_1_, …, *ν*_n_, e_n+1_, y) is a unitig. Fig. 3a illustrates an example of the de Bruijn graph and the relevant constructs defined.

**Figure 3:**
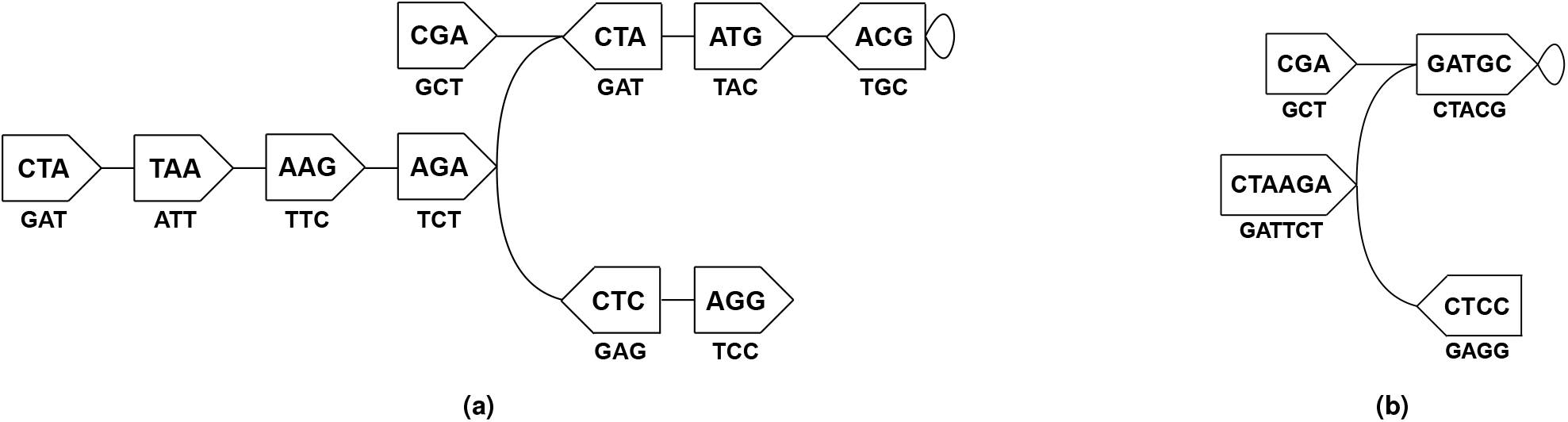
A (bidirected) edge-centric de Bruijn graph G(𝒮, k) for a set 𝒮 = {CTAAGAT, CGATGCA, TAAGAGG} of strings and k-mer size k = 3 in (a), and its compacted form G_c_(𝒮, k) in (b). In the graphs, the vertices are denoted with pentagons—the flat and the cusped ends depict the front and the back sides respectively, and each edge corresponds to some 4-mer(s) in s ∈ 𝒮. In (a), the vertices are the canonical forms of the k-mers in s ∈ 𝒮. The canonical string 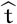 associated to each vertex *ν* is labelled inside *ν*, to be spelled in the direction from *ν*’s front to its back. Using 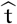, we also refer to *ν*. The label beneath *ν* is 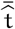, and is to be spelled in the opposite direction (i.e. back to front). For example, consider the 4-mer CGAT, an edge e in G(𝒮, 3). e connects the 3-mers x = pre (e) = CGA and y = suf (e) = GAT, the vertices being 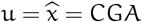 and 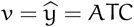 respectively. x is canonical and thus e exits through u ‘s back; whereas y is non-canonical and hence e enters t hrough *ν*’s back. (CTA, TAA, AAG) is a walk, a path, and also a unitig (edges not listed). (CGA, ATC, ATG) is a walk and a path, but not a unitig—the internal vertex ATC has multiple incident edges at its back. The unitig (CTA, TAA, AAG) is not maximal, as it can be extended farther through AAG’s back. Then it becomes maximal and spells CTAAGA. There are four such maximal unitigs in G(𝒮, 3), and contracting each into a single vertex produces G_c_(𝒮, 3), in (b). There are two different maximal path covers of G(𝒮, 3): spelling {CTAAGATGC, CGA, CCTC} and {CCTCTTAG, CGATGC}.

A *compacted de Bruijn graph* G_c_(𝒮, k) is obtained through collapsing each maximal unitig of the de Bruijn graph G(𝒮, k) into a single vertex, through contracting its constituent edges (75). Fig. 3b shows the compacted form of the graph in Fig. 3a. Given a set 𝒮 of strings and an integer k > 0, the problem of constructing the compacted de Bruijn graph G_c_(𝒮, k) is to compute the maximal unitigs of G(𝒮, k). ^7^

A *vertex-disjoint path cover* 𝒫 of G(𝒮, k) = (𝒱, ℰ) is a set of paths such that each vertex *ν* ∈ 𝒱 is present in exactly one path p ∈ 𝒫. Unless stated otherwise, we refer to this construct simply as path cover. A *maximal path cover* is a path cover 𝒫 such that no two paths in 𝒫 can be joined into one single path, i.e. there exists no p_1_, p_2_ ∈ 𝒫 such that p_1_ = (*ν*_0_, e_1_, …, e_n_, x), 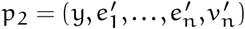, and there is some edge {(x, s_x_),(y, s_y_)} ∈ ℰ connecting the sides of x and y that are not incident to e_n_ and 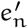, respectively. Fig. 3a provides examples of such.

### 3.3. Algorithm

Given a set ℛ, either of short-reads sequenced from some biological sample, or of reference sequences, the construction of the compacted de Bruijn graph G_c_(ℛ, k) for some k > 0 is a data reduction problem in computational genomics. A simple algorithm to construct the compacted graph G_c_ involves constructing the ordinary de Bruijn graph G(ℛ, k) at first, and then applying a graph traversal algorithm (76) to extract all the maximal non-branching paths in G. However, this approach requires constructing the ordinary graph and retaining it in memory for traversal (or organizing it in a way that it can be paged into memory for efficient traversal). Both of these tasks can be infeasible for large enough input samples. This prompts the requirement of methods to construct G_c_ directly from ℛ, by-passing G. Cuttlefish 2 is an asymptotically and practically efficient algorithm performing this task.

Another practical aspect of the problem is that the sequenced reads typically contain errors (77). This is offset through increasing the sequencing depth—even if a read r ∈ ℛ contains some erroneous symbol at index i of the underlying sequence being sampled, a high enough sequencing depth should ensure that some other reads in ℛ contain the correct nucleotide present at index i. Thus, in practice, these erroneous symbols need to be identified—usually heuristically—and the vertices and the edges of the graph corresponding to them need to be taken into account. Cuttlefish 2 discards the edges, i.e. (k+1)-mers, that each occur less than some threshold parameter f_0_, and only operates with the edges having frequencies ≥ f_0_.

#### 3.3.1. Implicit traversals over G(ℛ, k)

When the input is a set 𝒮 of references, the Cuttlefish algorithm (44) makes a complete graph traversal on the reference de Bruijn graph ^8^ G(𝒮, k) through a linear scan over each s ∈ 𝒮. Also, the concept of *sentinels* ^9^ in G(𝒮, k) ensures that a unitig can not span multiple input sequences. In one complete traversal, the branching vertices are characterized through obtaining a particular set of neighborhood relations; and then using another traversal, the maximal unitigs are extracted.

For a set ℛ of short-reads however, a linear scan over a read r ∈ ℛ may not provide a walk in G(ℛ, k), since r may contain errors, which break a contiguous walk. Additionally, the concept of sentinels is not applicable for reads. Therefore, unitigs may span multiple reads. For a unitig u spanning the reads r_1_ and r_2_ consecutively, it is not readily inferrable that r_2_ is to be scanned after r_1_ (or the reverse, for an oppositely-oriented traversal) while attempting to extract u directly from ℛ, as the reads are not linked in the input for this purpose. Hence, contrary to the reference-input algorithm (44) where complete graph traversal is possible with just |ℛ| different walks when ℛ consists of reference sequences, Cuttlefish 2 resorts to making implicit piecewise-traversals over G(ℛ, k).

For the purpose of determining the branching vertices, the piecewise-traversal is completely coarse—each walk traverses just one edge. Making such walks, Cuttlefish 2 retains just enough adjacency information for the vertices (i.e. only the internal edges of the unitigs) so that the unitigs can then be piecewise-constructed without the input ℛ. Avoiding the erroneous vertices is done through traversing only the *solid* edges ((k + 1)-mers occurring ≥ f_0_ times in ℛ, where f_0_ is a heuristically-set input parameter).

Stitching together the pieces of a unitig is accomplished by making another piecewise-traversal over G(ℛ, k), not by extracting those directly from the input sequences (opposed to Cuttlefish (44)). Each walk comprises the extent of a maximal unitig—the edges retained by the earlier traversal are used to make the walk and to stitch together the unitig.

In fact, the graph traversal strategy of Cuttlefish for reference inputs 𝒮 is a specific case of this generalized traversal, where a complete graph traversal is possible through a *linear* scan over the input, as each s ∈ 𝒮 constitutes a complete walk over G({s}, k). Besides, the maximal unitigs are tiled linearly in the sequences s ∈ 𝒮, and determining their terminal vertices is the core problem in that case; as extraction of the unitigs can then be performed through walking between the terminal vertices by scanning the ∈ 𝒮.

#### 3.3.2. A deterministic finite automaton model for vertices

While traversing the de Bruijn graph G(ℛ, k) = (𝒱, ℰ) for the purpose of determining the maximal unitigs, it is sufficient to only keep track of information for each side s_*ν*_ of each vertex *ν* ∈ 𝒱 that can categorize it into one of the following classes:

1. no edge has been observed to be incident to s_*ν*_ yet
2. s_*ν*_ has multiple distinct incident edges
3. s_*ν*_ has exactly one distinct incident edge, for which there are |Σ| = 4 possibilities (see Lemma 2, Suppl. Sec. 3).

Thus there are six classes to which each s_*ν*_ may belong, and since *ν* has two sides, it can be in one of 6 × 6 = 36 distinct configurations. Each such configuration is referred to as a *state*.

Cuttlefish 2 treats each vertex *ν* ∈ 𝒱 as a Deterministic Finite Automaton (DFA) M_*ν*_ = (𝒬, Σ′, δ, q_0_, 𝒬′):

##### States

𝒬 is the set of the possible 36 states for the automaton. Letting the number of distinct edges at the front with f and at the back with b for a vertex *ν* with a state q, and based on the incidence characteristics of *ν*, the states q ∈ 𝒬 can be partitioned into four disjoint *state-classes*: (1) *fuzzy-front fuzzy-back* (f ≠ 1, b ≠, (2) *fuzzy-front unique-back* (f ≠ 1, b = 1), (3) *unique-front fuzzy-back*, (f = 1, b ≠ 1), and (4) *unique-front unique-back* (f = 1, b = 1).

##### Input symbols

Σ′ = {(s, c) : s ∈ {front, back}, c ∈ Σ} is the set of the input symbols for the automaton. Each incident edge e of a vertex u is provided as input to u’s automaton. For u, an incident edge e = {(u, s_u_),(*ν*, s_*ν*_)} can be succinctly encoded by its incidence side s_u_ and a symbol c ∈ Σ —the (k + 1)-mer 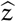 for e is one of the following, depending on s_u_ and whether 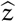 is exiting or entering u: u · c, ū · c, c · u, or c ·ū.

##### Transition function

δ : 𝒬 × Σ′→ 𝒬 \ {q_0_} is the function controlling the state-transitions of the automaton. Fig. 4 illustrates the state-transition diagram for an automaton as per δ.

**Figure 4:**
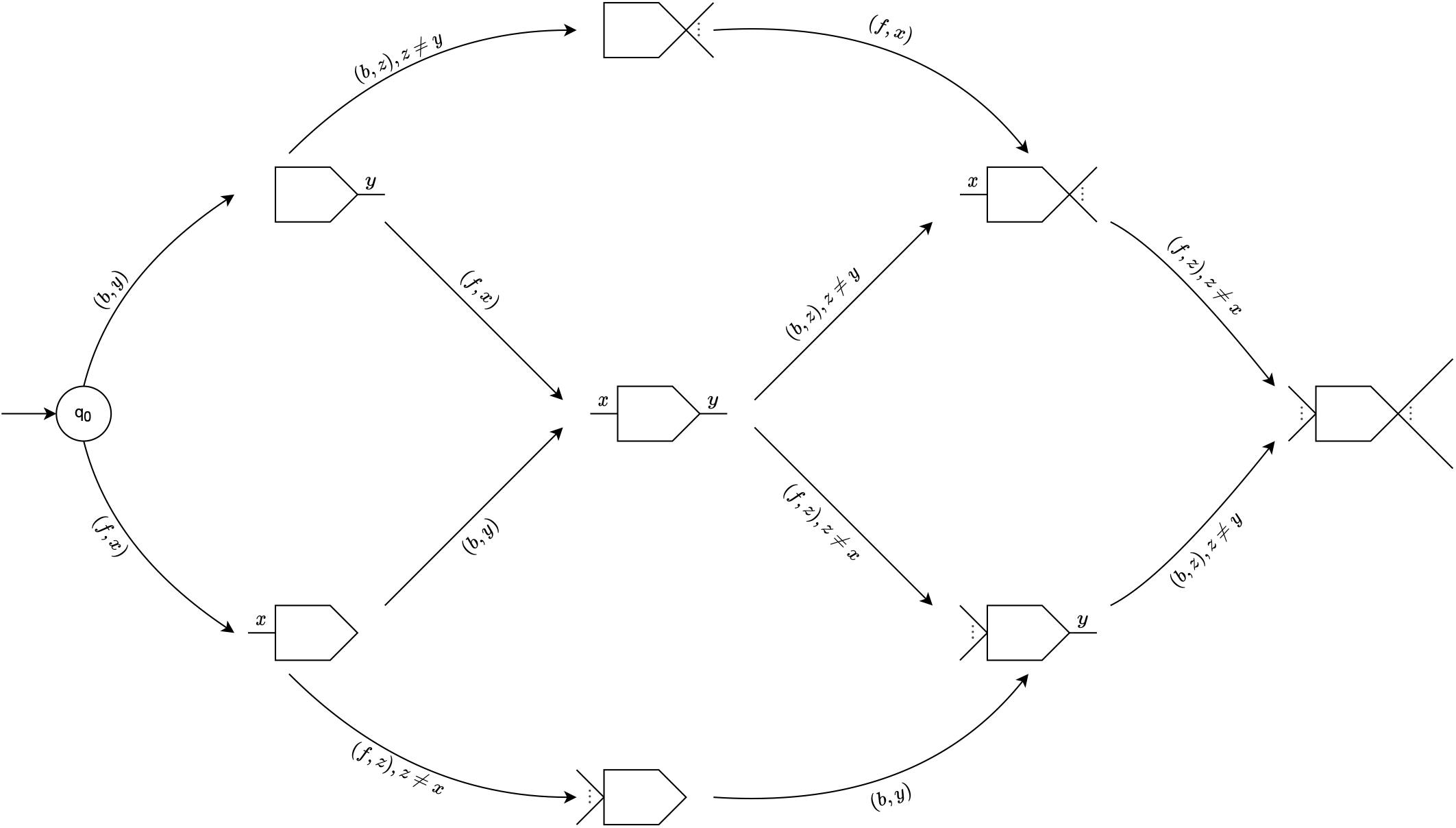
The state-transition diagram for an automaton M_*ν*_ = (𝒬, Σ′, δ, q_0_, 𝒬′). Each node in the diagram represents a collection of states q ∈ 𝒬, and q_0_ is the initial state of M_*ν*_. A node may represent multiple states collectively. For example, the node at the center of the figure with the symbols x and y at its flat and cusped ends respectively represents 16 states (all the ones from the state-class *unique-front unique-back*). Thus each node 𝒬_k_ represents a disjoint subset of 𝒬. The pictorial shape of 𝒬_k_ is similar to that of a de Bruijn graph vertex (see Fig. 3), and denotes the incidence characteristics of the vertices having their automata in states in 𝒬_k_: for a vertex *ν* with its automaton in state q_k_ ∈ 𝒬_k_, a unique symbol at side s of 𝒬_k_ means that *ν* has a distinct edge at side s, ellipsis means multiple unique edges, and absence of any symbol means no edge has been observed incident to that side. A directed edge (𝒬_i_, 𝒬_j_) labelled with (s, c) denotes transitions from a state q_i_ ∈ 𝒬_i_ to a state q_j_ ∈ 𝒬_j_, and (s, c) symbolizes the corresponding input fed to an automaton at state q_i_ for that transition to happen. That is, δ q_i_,(s, c) = q_j_. Thus these edges pictorially encode the transition function δ. For the automaton M_*ν*_ of a vertex *ν*, an input (s, c) means that an edge e is being added to its side s ∈ {f, b}; along with s and *ν*, the character c ∈ Σ succinctly encodes e. f and b are shorthands for front and back, respectively. Self-transition is possible for each state q ∈ 𝒬′, and are not shown here for brevity.

##### Initial state

q_0_ ∈ 𝒬 is the initial state of the automaton. This state corresponds to the configuration of the associated vertex at which no edge has been observed to be incident to either of its sides.

##### Accept states

𝒬′ = 𝒬 \ {q_0_} is the set of the states corresponding to vertex-configurations having at least one edge. ^10^

#### 3.3.3. Algorithm overview

We provide here a high-level overview of the Cuttlefish 2(ℛ, k, f_0_) algorithm. The input to the algorithm is a set ℛ of strings, an odd integer k > 0, and an abundance threshold f_0_ > 0; the output is the set of strings spelled by the maximal unitigs of the de Bruijn graph G(ℛ, k).

Cuttlefish 2(ℛ, k, f_0_)

1. ℰ ← Enumerate-Edges(ℛ, k, f_0_)
2. 𝒱 ← Extract-Vertices(ℰ)
3. h ←Construct-Minimal-Perfect-Hash(𝒱)
4. S ← Compute-Automaton-States(ℰ, h)
5. 𝒰 ← Extract-Maximal-Unitigs(𝒱, h, S)

Cuttlefish 2(ℛ, k, f_0_) executes in five high-level stages, and Fig. 1 illustrates these steps. Firstly, it enumerates the set ℰ of edges, i.e. (k + 1)-mers that appear at least f_0_ times in ℛ. Then the set 𝒱 of vertices, i.e. k-mers are extracted from ℰ. Having the distinct k-mers, it constructs a minimal perfect hash function h over 𝒱. At this point, a hash table structure is set up for the automata—the hash function being h, and each hash bucket having enough bits to store a state encoding. Then, making a piecewise traversal over G(ℛ, k) using ℰ, Cuttlefish 2 observes all the adjacency relations in the graph, making appropriate state transitions along the way for the automata of the vertices u and *ν* for each edge { (u, s_u_),(*ν*, s_*ν*_)}. After all the edges in ℰ are processed, each automaton M_*ν*_ resides in its correct state. Due to the design characteristic of the state-space 𝒬 of M_*ν*_, the internal vertices of the unitigs in G(ℛ, k), as well as the non-branching sides of the branching vertices have their incident edges encoded in their states. Cuttlefish 2 retrieves these unitig-internal edges and stitches them together in chains until branching vertices are encountered, thus extracting the maximal unitigs implicitly through another piecewise traversal, with each walk spanning a maximal unitig.

These major steps in the algorithm are detailed in the following subsections. Then we analyze the asymptotic characteristics of the algorithm in Sec. 3.4. Finally, we provide a proof of its correctness in Theorem 1 (see Suppl. Sec. 3).

#### 3.3.4. Edge set construction

The initial enumeration of the edges, i.e. (k + 1)-mers from the input set ℛ is performed with the KMC 3 algorithm (72). KMC 3 enumerates the 𝓁-mers of its input in two major steps. Firstly, it partitions the 𝓁-mers based on *signatures*—a restrictive variant of *minimizers* ^11^. Maximal substrings from the input strings with all their 𝓁-mers having the same signature (referred to as *super* 𝓁*-mers*) are partitioned into bins corresponding to the signatures. Typically the number of bins is much smaller than the number of possible signatures, and hence each bin may contain strings from multiple signatures (set heuristically to balance the bins). In the second phase, for each partition, its strings are split into substrings called (𝓁, x)*-mers*—an extension of 𝓁-mers. These substrings are then sorted using a most-significant-digit radix sort (78) to unify the repeated 𝓁-mers in the partition. For 𝓁 = k + 1, the collection of these deduplicated partitions constitute the edge set ℰ.

#### 3.3.5. Vertex set extraction

Cuttlefish 2 extracts the distinct canonical k-mers—vertices of G(ℛ, k)—from ℰ through an extension of KMC 3 (72) (See Suppl. Sec. 2.1). For such, taking ℰ as input, KMC 3 treats each (k + 1)-mer e ∈ ℰ as an input string, and enumerates their constituent k-mers applying its original algorithm. Using ℰ instead of ℛ as input reduces the amount of work performed in this phase through utilizing the work done in the earlier phase— skipping another pass over the entire input set ℛ, which can be computationally substantial.

#### 3.3.6. Hash table structure setup

An associative data structure is required to store the transitioning states of the automata of the vertices of G(ℛ, k). That is, association of some encoding of the states to each canonical k-mer is required. Some off-the-shelf hash table can be employed for this purpose. Due to hash collisions, general-purpose hash tables typically need to store the keys along with their associated data—the key set 𝒱 may end up taking klog_2_ |Σ| = 2k bits/k-mer in the hash table ^12^. In designing a more efficient hash table structure, Cuttlefish 2 makes use of the facts that: (i) the set 𝒱 of keys is static—no alien keys will be encountered while traversing the edges in ℰ, since 𝒱 is constructed from ℰ; and (ii) 𝒱 has been enumerated at this point. A *Minimal Perfect Hash Function* (MPHF) is applicable in this setting. Given a set 𝒦 of keys, a *perfect hash function* over 𝒦 is an injective function h : 𝒦 → ℤ, i.e. ∀ k_1_, k_2_ ∈ 𝒦, k_1_ ≠ k_2_ ⇔ h(k_1_) ≠ h(k_2_). h is minimal when its image is [0, |𝒦|), i.e. an MPHF is an injective function h : 𝒦 → [0, |𝒦|). By definition, an MPHF does not incur collisions. Therefore when used as the hash function for a hash table, it obviates the requirement to store the keys with the table structure. Instead, some encoding of the MPHF needs to be stored in the structure.

To associate the automata to their states, Cuttlefish 2 uses a hash table with an MPHF as the hash function. An MPHF h over the vertex set 𝒱 is constructed with the BBHash algorithm (79). For the key set 𝒱_0_ = 𝒱, BBHash constructs h through a number of iterations. It maps each key *ν* ∈ 𝒱_0_ to [ 1, γ|𝒱_0_| ] with a classical hash function h_0_, for a provided parameter γ > 0. The collision-free hashes in the hash codomain [ 1, γ|𝒱_0_| ] are marked in a bit-array A_0_ of length γ|𝒱_0_|. The colliding keys are collected into a set 𝒱_1_, and the algorithm is iteratively applied on 𝒱_1_. The iterations are repeated until either some 𝒱_n_ is found empty, or a maximum level is reached. The bit-arrays A_i_ for the iterations are concatenated into an array A, which along with some other meta-data, encode h. A has an expected size of γe^1/γ^|𝒱| bits (79). γ trades off the encoding size of h with its computation time. γ = 2 provides a reasonable trade-off, with the size of h being ≈ 3.7 bits/vertex. ^13^ Note that, the size is independent of k, i.e. the size of the keys.

For the collection of hash buckets, Cuttlefish 2 uses a linear array (81) of size |𝒱|. Since each bucket is to contain some state q ∈ 𝒬, ⌈log_2_ |𝒬|⌉ = ⌈log_2_ 36⌉ = 6 bits are necessary (and also sufficient) to encode q. Therefore Cuttlefish 2 uses 6 bits for each bucket. The hash table structure is thus composed of an MPHF h and a linear array S: for a vertex *ν*, its (transitioning) state q_*ν*_ is encoded at the index h(*ν*) of S, and in total the structure uses ≈ 9.7 bits/vertex.

#### 3.3.7. Automaton states computation

Given the set ℰ of edges of a de Bruijn graph G(ℛ, k) and an MPHF h over its vertex set 𝒱, the Compute-Automaton-States(ℰ, h) algorithm computes the state of the automaton M_*ν*_ of each *ν* ∈ 𝒱.

It initializes each automaton M_w_ with q_0_—the initial state corresponding to no incident edges. Then for each edge

Compute-Automaton-States(ℰ, h)

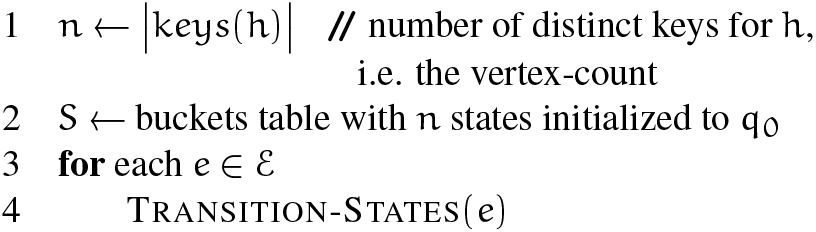

Transition-States(e)

1. u → pre_k_ (e), *ν* → suf_k_ (e)
2. s_u_ → Exit-side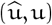
3. s_*ν*_ → Entrance-Side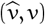
4. *c*_u_→ e_k+1_ if s_u_ = back,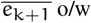
5. *c*_*u*_→ e_1_ if s_*ν*_ = front,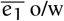
6. 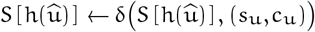
7. 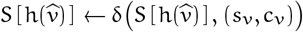

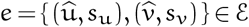, connecting the vertex 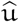 via its side s_u_ to the vertex 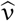 via its side s_*ν*_, it makes appropriate state transitions for M_u_ and M_*ν*_, the automata of 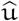 and 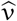 respectively. For each endpoint 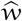 of e, (s_w_, c_w_) is fed to M_w_, where c_w_ ∈ Σ. Together with 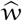, s_w_ and c_w_ encode e. The setting policy for c_w_ is described in the following. Technicalities relating to loops are accounted for in the Cuttlefish 2 implementation, but are omitted from discussion for simplicity.

e has two associated (k + 1)-mers: z and 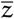. Say that z =u ⊙^k −1^ *ν*. Based on whether 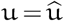 holds or not, e is incident to either 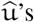 back or front. As defined (see Sec. 3.2), if it is incident to the back, then 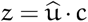 otherwise, 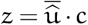, where c = e_k+1_. In these cases respectively, 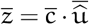, and 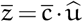. For consistency, Cuttlefish 2 always uses a fixed form of e for 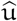—either z or 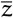—to provide it as input to M_u_: the one containing the k-mer u in its canonical form. So if e is at 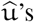 back, the 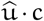 form is used for e, and (back, c) is fed to M_u_ ; otherwise, e is expressed as 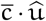 and (front, 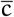) is the input for M_u_. The encoding (s_v_, c^′^) of e for 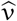 is set similarly.

#### 3.3.8. Maximal Unitigs Extraction

Given the set 𝒱 of vertices of a de Bruijn graph G(ℛ, k), an MPHF h over 𝒱, and the states-table S for the automata of *ν* ∈ 𝒱, the Extract-Maximal-Unitigs(𝒱, h, S) algorithm assembles all the maximal unitigs of G(ℛ, k).

The algorithm iterates over the vertices in 𝒱. For some vertex 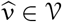, let p be the maximal unitig containing 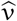. p can be broken into two subpaths: p_b_ and p_f_, overlapping only at 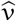. The Extract-Maximal-Unitigs(𝒱, h, S) algorithm extracts these subpaths separately, and joins them at 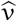 to construct p. Then p’s constituent vertices are marked by transitioning their automata to some special states (not shown in Fig. 4), so that p is extracted only once.

The subpaths p_b_ and p_f_ are extracted by initiating two walks: one from each of 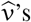 sides back and front, using the Walk-Maximal-Unitig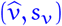 algorithm. Each walk continues on until a *flanking vertex* 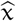 is encountered. For a vertex 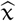, let q_x_ denote the state of 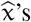 automaton and 𝒞_x_ denote q_x_’s

Extract-Maximal-Unitigs(𝒱, h, S)

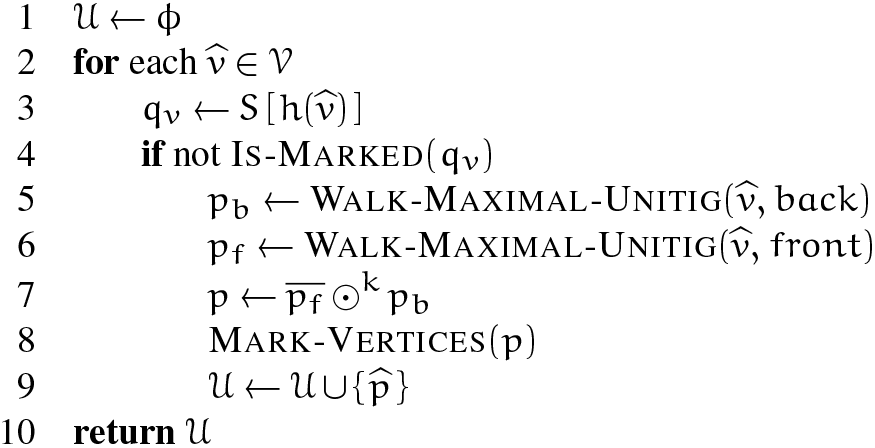

Walk-Maximal-Unitig(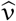,sv)

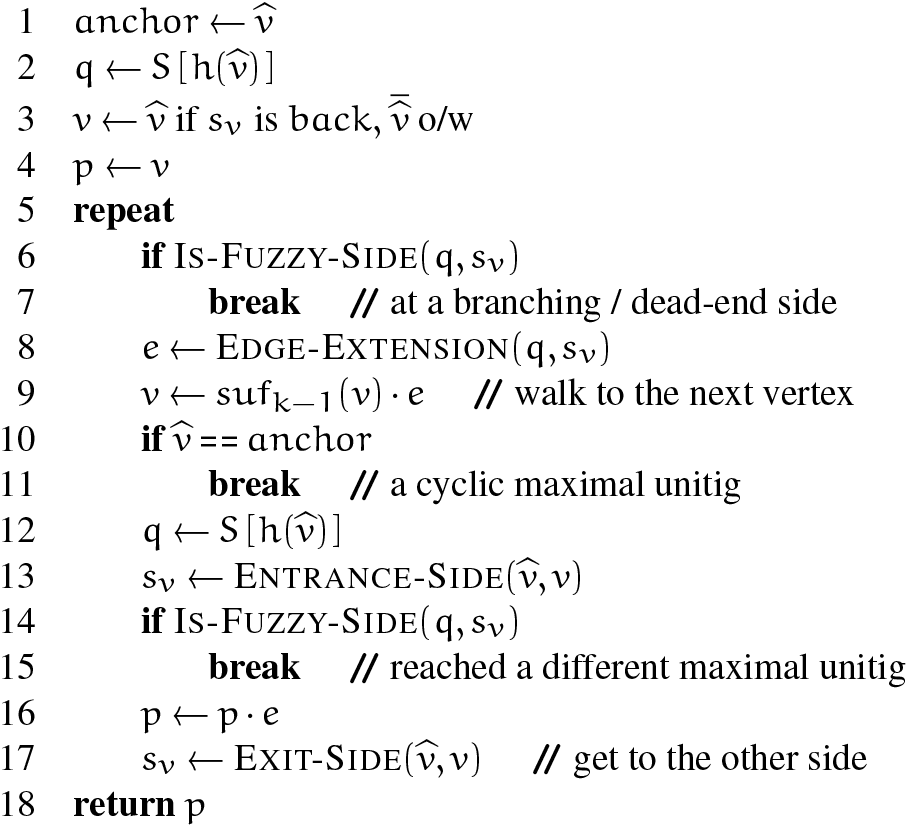

state-class. Then 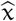 is noted to be a flanking vertex iff:

1. either 𝒞_x_ is not *unique-front unique-back*;
2. or 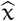 connects to the side s of a vertex 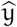 such that:
  a. 𝒞_y_ is *fuzzy-front fuzzy-back*; or
  b. s_y_ = front and 𝒞_y_ is *fuzzy-front unique-back*; or
  c. s_y_ = back and 𝒞_y_ is *unique-front fuzzy-back*.

Lemma 3 (see Suppl. Sec. 3) shows that the flanking vertices in G(ℛ, k) are precisely the endpoints of its maximal unitigs. The Walk-Maximal-Unitig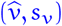 algorithm initiates a walk w from 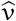, exiting through its side s_v_. It fetches 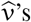 state q_v_ from the hash table. If q_v_ is found to be not belonging to the state-class *unique-front unique-back* due to s_v_ having ≠1 incident edges, then 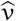 is a flanking vertex of its containing maximal unitig p, and p has no edges at s_v_. Hence w terminates at 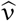. Otherwise, s_v_ has exactly one incident edge. The walk algorithm makes use of the fact that, the vertex-sides s_u_ that are internal to the maximal unitigs in G(ℛ, k) contain their adjacency information encoded in the states q_u_ of their vertices 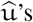 automata, once the Compute-Automaton-States(ℰ, h) algorithm is executed. Thus, it decodes q_v_ to get the unique edge 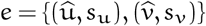 incident to s_v_. Through e reaches the neighboring vertex 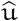, at its side 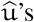 state q_u_ is fetched, and if q_u_ is found not to be in the class *unique-front unique-back* due to s_u_ having incident edges, then both 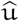 and 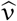 are flanking vertices (for different maximal unitigs), and w retracts to and stops at 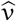. Otherwise, e is internal to p, and w switches to the other side of 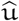, proceeding on similarly. It continues through vertices 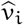 in this manner until a flanking vertex of p is reached, stitching together the edges along the way to construct a sub-path of p.

A few constant-time supplementary procedures are used throughout the algorithm. Is-Fuzzy-Side(q, s) determines whether a vertex with the state q has 0 or > 1 edges at its side s. Edge-Extension(q, s) returns an encoding of the edge incident to the side s of a vertex with state q. Entrance-Side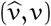 and Exit-Side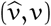 returns the side used to enter (and exit) the vertex 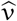 when its k-mer form *ν* is observed.

#### 3.3.9. Maximal path-cover extraction

We discuss here how Cuttlefish 2 might be modified so that it can extract a maximal path cover of a de Bruijn graph G(ℛ, k). For such, only the Compute-Automaton-States step needs to be modified, and the rest of the algorithm remains the same. Given the edge set ℰ of the graph G(ℛ, k) and an MPHF h over its vertex set 𝒱, Compute-Automaton-States-Path-Cover(ℰ,h) presents the modified DFA states computation algorithm.

The maximal path cover extraction variant of Cuttlefish 2 works as follows. It starts with a trivial path cover 𝒫 _0_ of G(ℛ, k): each *ν* ∈ 𝒱 constitutes a single path, spanning the subgraph G^′^ (𝒱, ∅). Then it iterates over the edges e ∈ ℰ (with |ℰ| = m) in arbitrary order. We will use 𝒫_i_ to refer to the path cover after having visited i edges. At any given point of the execution, the algorithm maintains the invariant that 𝒫_i_ is a maximal path cover of the graph G^′^(𝒱, ℰ^′^), where ℰ^′^ ⊆ ℰ (with |ℰ^′^| = i) is the set of the edges examined until that point. When examining the (i + 1)’th edge e = {(u, s_u_),(*ν*, s_v_)}, it checks whether e connects two different paths in 𝒫_i_ into one single path: this is possible iff the sides s_u_ and s_v_ do not have any incident edges already in ℰ^′^, i.e. the sides are empty in G^′^(𝒱, ℰ^′^). If this is the case, the paths are joined in 𝒫_i+1_ into a single path containing the new edge e. Otherwise, the path cover remains unchanged so that 𝒫_i+1_ = 𝒫_i_. By definition, 𝒫_i+1_ is a path cover of G^′^(𝒱, ℰ^′^ ∪ {e}), as e could only affect the paths (at most two) in 𝒫_i_ containing u and *ν*, while the rest are unaffected and retain maximality—thus the invariant is maintained. By induction, 𝒫_m_ is a path cover of G(𝒱, ℰ) once all the edges have been examined, i.e. when ℰ^′^ = ℰ.

By making state transitions for the automata only for the edges present internal to the paths p ∈ 𝒫_m_, the Compute-Automaton-States-Path-Cover(ℰ,h) algorithm seamlessly captures the subgraph 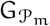 of G(ℛ, k) that is induced by the path cover 𝒫_m_. 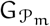 consists of a collection of disconnected maximal paths, and thus there exists no branching in 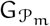. Consequently, each of these maximal paths is a maximal unitig of 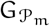. The sub-sequent Extract-Maximal-Unitigs algorithm operates using the DFA states collection S computed at this step, and therefore it extracts precisely these maximal paths.

Compute-Automaton-States-Path-Cover(ℰ, h)

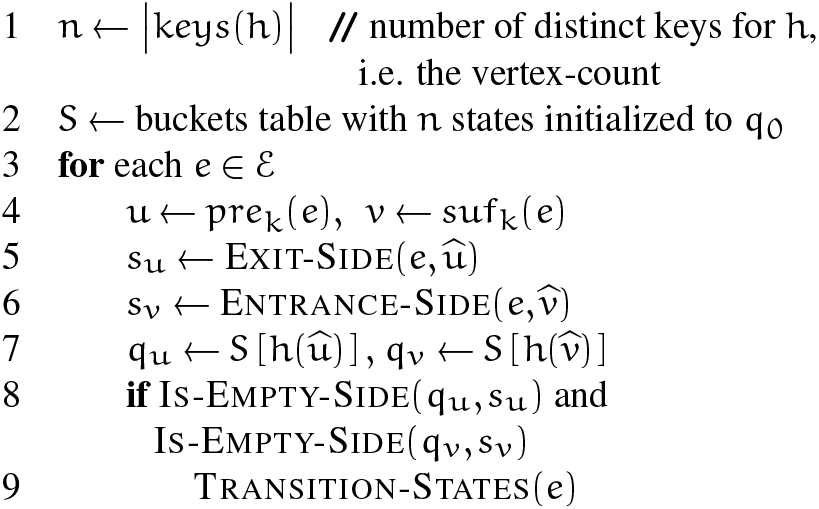

#### 3.3.10. Parallelization

Cuttlefish 2 is highly parallelizable on a shared-memory multi-core machine. The Enumerate-Edges and the Extract-Vertices steps, using KMC 3 (72), are parallelized in their constituent phases via parallel distribution of the input (k+ 1)-mers (and k-mers) into partitions, and sorting multiple partitions in parallel.

The Compute-Minimal-Perfect-Hash step using BB-Hash (79) parallelizes the construction through distributing disjoint subsets 𝒱_i_ of the vertices to the processor-threads, and the threads process the 𝒱_i_’s in parallel.

The next two steps, Compute-Automaton-States and Extract-Maximal-Unitigs, both (piecewise) traverse the graph through iterating over ℰ and 𝒱 respectively. The processor-threads are provided disjoint subsets of ℰ and 𝒱 to process in parallel. Although the threads process different edges in Compute-Automaton-States, multiple threads may access the same automaton into the hash table simultaneously, due to edges sharing endpoints. Similarly in Extract-Maximal-Unitigs, though the threads examine disjoint vertex sets, multiple threads simultaneously constructing the same maximal unitig from its different constituent vertices can access the same automaton concurrently, at the walks’ meeting vertex. Cuttlefish 2 maintains exclusive access to a vertex to one thread at a time through a sparse set ℒ of locks. Each lock l ∈ ℒ guards a disjoint set 𝒱_i_ of vertices, roughly of equal size. With p processor-threads and assuming all p threads accessing the hash table at the same time, the probability of two threads concurrently probing the same lock at the same turn is 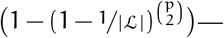 this is minuscule with an adequate | ℒ|. ^14^

### 3.4. Asymptotics

In this section, we analyze the computational complexity of the Cuttlefish 2(ℛ, k, f_0_) algorithm when executed on a set ℛ of strings, given a k value, and a threshold factor f_0_ for the edges in G(ℛ, k). ℰ is the set of the (k + 1)-mers occurring ≥ f_0_ times in ℛ, and 𝒱 is the set of the k-mers in ℰ. Let 𝓁 be the total length of the strings r ∈ ℛ, n be the vertex-count |𝒱|, and m be the edge-count |ℰ|.

#### 3.4.1. Time complexity

Cuttlefish 2 represents j-mers with 64-bit machine-words—packing 32 symbols into a single word. Let w_j_ denote the number of words in a j-mer, i.e. 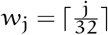

Note that the number of (k + 1)-mers in ℛ is upper-bounded by 𝓁. The Enumerate-Edges step uses the KMC 3 (72) algorithm. At first, it partitions the (k + 1)-mers into buckets based on their signatures. With a rolling computation, determining the signature of a (k + 1)-mer takes an amortized constant time. Assigning a (k + 1)-mer to its bucket then takes time 𝒪(w_k+1_), and the complete distribution takes 𝒪 (w_k+1_𝓁). ^15^ As each (k + 1)-mer consists of w_k+1_ words, radix-sorting a bucket of size B_i_ takes time 𝒪(B_i_w_k+1_). So in the second step, for a total of b buckets for ℛ, the cumulative sorting time is 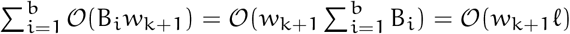 Thus Enumerate-Edges takes time 𝒪(𝓁w_k+1_).

The Extract-Vertices step applies KMC 3 (72) with ℰ as input, and hence we perform a similar analysis as earlier. Each e ∈ ℰ comprises two k-mers. So partitioning the k-mers takes time 𝒪(2mw_k_), and radix-sorting the buckets takes 𝒪(w_k_ Σ B_i_) =𝒪(2mw_k_). Therefore Extract-Vertices takes time 𝒪(mw_k_).

The Construct-Minimal-Perfect-Hash step applies the BBHash (79) algorithm to construct an MPHF h over 𝒱. It is a multi-pass algorithm—each pass i tries to assign final hash values to a subset 𝒦_i_ of keys. Making a bounded number of passes over sets 𝒦_i_ of keys—shrinking in size— it applies some classical hash h_i_ on the x ∈ 𝒦_i_ in each pass. For some x ∈ 𝒦_i_, iff h_i_(x) is free of hash collisions, then x is not propagated to 𝒦_i+1_. Provided that the h_i_’s are uniform and random, each key *ν* ∈ 𝒱 is hashed with the h_i_’s an expected 𝒪(e^1/ ϒ^) times (79), an exponentially decaying function on the γ parameter. Given that h_i_’s are constant time on machine-words, computing h_i_(*ν*) takes time 𝒪(w_k_). Then the expected time to assign its final hash value to a *ν* ∈ 𝒱 is H(k) = 𝒪(w_k_e^1/ϒ^). Therefore Construct-Minimal-Perfect-Hash takes an expected time 𝒪(nH(k)). Note that, querying h, i.e. computing h(*ν*) also takes time H(k), as the query algorithm is multi-pass and similar to the construction.

The Compute-Automaton-States step initializes the n automata with the state q_0_, taking time 𝒪(n). Then for each edge e ∈ ℰ, it fetches its two endpoints’ states from the hash table in time 2H(k), updating them if required. In total there are 2m hash accesses, and thus Compute-Automaton-States takes time 𝒪(n+ mH(k)).

The Extract-Maximal-Unitigs step scans through each vertex *ν* ∈ 𝒱, and walks the entire maximal unitig p containing v. The state of each vertex in p is decoded to complete the walk—requiring |p| hash table accesses, taking time |p|H(k). If the flanking vertices of p are non-branching, then the walk also visits their neighboring vertices that are absent in p, at most once per each endpoint. Once extracted, all the vertices in p are marked so that p is not extracted again later on—this takes time |p|H(k), and can actually be done in time 𝒪(|p|) by saving the hash values of the path vertices while constructing p. Thus traversing all the u_i_’s in the maximal unitigs set 𝒰 takes time 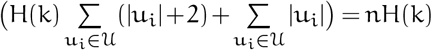.

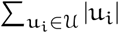 equates to n because the set of the maximal unitigs 𝒰 forms a vertex decomposition of G(ℛ, k) (47). Thus Extract-Maximal-Unitigs takes time 𝒪(nH(k)).

In the brief analysis for the last three steps, we do not include the time to read the edges (𝒪(mw_k + 1_)) and the vertices (𝒪(nw_k_)) into memory, as they are subsumed by other terms.

Thus, Cuttlefish 2(ℛ, k, f_0_) has an expected running time 𝒪(𝓁w_k+1_ + mw_k_ + (n+ m)H(k), where 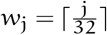, and ϒ > 0 is a constant. It is evident that the bottleneck is the initial Enumerate-Edges step, and it asymptotically subsumes the running time.

#### 3.4.2. Space complexity

Here, we analyze the working memory (i.e. RAM) required by the Cuttlefish 2 algorithm. The Enumerate-Edges step with KMC 3 (72) can work within a bounded memory space. Its partitioning phase distributes input k-mers into disk bins, and the k-mers are kept in working memory within a total space limit S, before flushes to disk. Radix-sorting the bins are done through loading bins into memory with sizes within S, and larger bins are broken into sub-bins to facilitate bounded-memory sort. As we discuss below, the graph traversal steps require a fixed amount of memory, determined linearly by n. As n is not computed until the completion of Extract-Vertices, we approximate it within the KMC 3 algorithm (see Suppl. Sec. 2.1), and then bound the memory for the KMC 3 execution appropriately. The next step of Extract-Vertices is also performed similarly within the same memory-bound.

The Construct-Minimal-Perfect-Hash step with BBHash (79) processes the key set 𝒱 in fixed-sized chunks. Each pass i with key set 𝒱_i_ has a bit-array A_i_ to mark h_i_(v) for all the *ν* ∈ 𝒱_i_, along with an additional bit-array C_i_ to detect the hash collisions. Both A_i_ and C_i_ have the size ϒ|𝒱_i_|. The finally concatenated A_i_’s is the output data structure A for the algorithm, and some C_i_ is present only during the pass i. A has an expected size of ϒe^1/ϒ^ n bits (79). |C_0_| = γ|𝒱_0_| = ϒn, and this is the largest collision array in the algorithm’s lifetime. Thus, an expected loose upper-bound of the memory usage in this step is 𝒪(|A| + |C_0_| = 𝒪 ((e^1/ϒ^ + 1) ϒn) bits.

At this point in the algorithm, a hash table structure is set up for the automata. Together, the hash function h and the hash buckets collection S take an expected space of (ϒe^1/ϒ^n + n ⌈log2| 𝒬 | ⌉) = (ϒe^1/ϒ^ + 6)n bits.

The Compute-Automaton-States step scans the edges in ℰ in fixed-sized chunks. For each e ∈ ℰ, it queries and updates the hash table for the endpoints of e as required. Similarly, the Extract-Maximal-Unitigs step scans the vertices in 𝒱 in fixed-sized chunks, and spells the containing maximal unitig of some *ν* ∈ 𝒱 through successively querying the hash table for the path vertices. The spelled paths are dumped to disk at a certain cumulative threshold size. Thus the only non-trivial memory usage by these steps is from the hash table. Therefore these graph traversal steps use ((ϒe^1/ϒ^ + 6)n+ 𝒪 (1) bits.

When ϒ ≤ 6, the hash table (i.e. the hash function and the bucket collection) is the the dominant factor in the algorithm’s memory usage, and Cuttlefish 2(ℛ, k, f_0_) consumes expected space 𝒪 ((ϒe^1/ϒ^ + 6)n). If γ > 6 is set, then it could be possible for the hash function construction memory to dominate. In practice, we adopt ϒ = 2, and the observed memory usage is ≈ 9.7n bits, translating to ≈ 1.2 bytes per distinct k-mer.

## Footnotes

1 From our observations, the distributions of k-mer frequencies and of (k+ 1)-mer frequencies on real data tend to agree closely, resulting in the same f_0_ for these experiments for both Cuttlefish 2 and the rest of the algorithms, as per the setting-policy used.

2 k-mers occurring frequently enough in input NGS reads are said to be solid k-mers, and the other ones are said to be weak (65).

3 Length 𝓁 of the longest contig such that all the contigs having lengths ≥ 𝓁 sum in size to at least 50% of the sum size of the contigs.

4 Analogous to N50, except for: (1) breaking the contigs into their constituent blocks that can be aligned to an associated reference sequence, and (2) replacing the sum size of contigs with the reference length.

5 As per this definition, 𝒱_2_ = 𝒦. We describe in Sec. 3.3 a practical consideration that implies that 𝒱_2_ need not necessarily be the same as 𝒦 when some *filtering* is applied on the input 𝒮 to generate 𝒦_+1_.

6 This is to account for the DNA being double-stranded, and a genomic string may come from either of these oppositely-oriented complementary strands.

7 Discarding orientations: the two unitigs (*ν*_0_, …, *ν*_n_) and (*ν*_n_, …, *ν*_0_) are topologically the same.

8 Introduced by Khan and Patro (44), based on the input to the de Bruijn graph constructions being either reference sequences or sequencing reads, the graphs are distinguished as either *reference* or *read de Bruijn graphs*.

9 A vertex *ν* is a sentinel if the first or the last k-mer x of an input string corresponds to *ν*. Let *ν*’s empty side in x be s_*ν*_. The graph G(𝒮, k) is modified such that s_*ν*_ connects to a special branching vertex ϒ—preventing unitigs containing *ν* to span farther through s_*ν*_.

10 Formally, 𝒬′ is the set of states reachable from q_0_ through transitions as per some definite patterns of input symbols. For our purposes, recognizing specific input patterns is not a concern—rendering this parameter redundant—we define it as the set of the final states an automaton can be in having fed all its inputs.

11 For a given j < 𝓁, a j-minimizer of an 𝓁-mer x is the smallest j-mer substring of x according to some specified function.

12 This can be improved by having 4^p^ different hash tables for 𝒱, for a fixed prefix length p ≤ k. Each hash table then accounts for keys of length (k − p).

13 It can be as low as ≈ 3 bits/vertex with γ = 1, at the expense of slower hashing. The theoretical lower limit for MPHFs is ≈ 1.44 bits/key (80).

14 The optimal (in regard to probability) value |ℒ| = |𝒱| is not used due to the locks’ memory usage.

15 This bound is not tight, as KMC 3 actually distributes sequences longer than (k + 1)-mers—reducing computation (see Sec. 3.3.4).

