## Supplementary material for "Scalable, ultra-fast, and low-memory construction of compacted de Bruijn graphs with Cuttlefish 2"

### 1. Results

**1.1. Choice of frequency thresholds.** The frequency threshold $f_0$ of $k$-mers ($(k+1)$-mers in case of CUTTLEFISH 2) for the algorithms when working with sequencing data was approximated so as to roughly minimize the misclassification rates of weak and solid $k$-mers in these experiments. This was performed based on approximate frequency distributions of the $k$-mer frequencies themselves, computed using the NTCARD tool (1). The heuristic setting-policy of $f_0$ is inspired from observations by Zhao et al. (2): the frequency distribution of erroneous $k$-mers tend to diminish exponentially, whereas that of error-free $k$-mers typically follow a normal distribution; and the intersecting point of these density functions can be a *reasonable* choice for $f_0$, which we approximated with NTCARD (1). Suppl. Fig. 1 shows some of these approximate distributions.

**1.2. Compacted graph construction for sequencing data.** Suppl. Table 1 contains the performance results of the evaluated tools for compacted de Bruijn graph construction from sequencing data.

Suppl. Table 2 shows the performance results on the human read set with a frequency cutoff of $f_0 = 2$.

**1.3. Compacted graph construction for reference collections.** Suppl. Table 3 shows the performance results of the evaluated tools for compacted de Bruijn graph construction from reference sequence collections.

**1.4. Timing-profile without $(k+1)$-mer (or $k$-mer) enumeration.** We tested the hypothesis of whether having a uniform $k$-mer enumerator for CUTTLEFISH 2 and BCALM 2 might significantly impact their performance difference. Suppl. Table 4 demonstrates the timing-profile of CUTTLEFISH 2 compared to BCALM 2, excluding their similar initial stage: $(k+1)$-mer and $k$-mer enumeration, respectively.

We find that CUTTLEFISH 2 still largely outperforms BCALM 2 in time. As for memory advantage, the BCALM 2 implementation has a "maximum memory" parameter, `-max-memory`, that could be used to restrict its memory-usage to the given argument value. In all our experiments, we set the value of `-max-memory` to the memory-usage incurred by CUTTLEFISH 2. But BCALM 2 did not strictly adhere to these limits: in both its $k$-mer counting and the subsequent compaction steps. [1] So replacing its $k$-mer enumeration step, which uses DSK (3), with KMC 3 would not necessarily constrain its memory-usage to the ones observed for CUTTLEFISH 2.

**1.5. Validation of the compacted de Bruijn graphs.** The validation of a compacted de Bruijn graph consists of checking three aspects of the graph: (1) completeness: whether the set of maximal unitigs contain all the $k$-mers from the original de Bruijn graph; (2) maximality: whether the output unitigs are actually maximal. and (3) branch-freeness: that the complete, maximal cover of the de Bruijn graph contains no paths having internal vertices than branch in the underlying de Bruijn graph.

Theoretically, the CUTTLEFISH 2 algorithm obtains all three of these criteria. Completeness is obtained trivially, and maximality and branch-freeness are obtained as per Theorem 1. We cross-checked the correctness of the actual implementation by validating the output graphs of CUTTLEFISH 2 against those of BCALM 2 and BIFROST. In doing so, we observed some small-scale differences in both the $k$-mer-content (completeness) and unitig-content (maximality). We provide some informal reasoning for these differences here:

***Vertex-centric versus Edge-centric de Bruijn graphs.*** For a given dataset, let its $k$-mer set be $\mathcal{V}$ and $(k+1)$-mer set be $\mathcal{E}$. Consider its vertex-centric de Bruijn graph $G_{\mathcal{V}}$ and its edge-centric de Bruijn graph $G_{\mathcal{E}}$. For ease of exposition, assume that there is no $k$-mer (or $(k+1)$-mer) filtering performed, and that the graphs do not have tips [2], which consist of a negligible number of vertices for real datasets. Both the graphs have the same vertex set $\mathcal{V}$.

By definition, any edge $e_1$ in $G_{\mathcal{E}}$ must also exist in $G_{\mathcal{V}}$. But this does not necessarily hold true in the opposite direction: some edge $e' = \{u, v\}$ in $G_{\mathcal{V}}$ could be absent in $G_{\mathcal{E}}$—although there exists a $(k-1)$-length overlap between the $k$-mers $u$ and $v$, the $(k+1)$-mer $u \odot^{k-1} v$ could be absent in $\mathcal{E}$. [3] Thus $G_{\mathcal{V}}$ must always have an equal or greater number of edges than $G_{\mathcal{E}}$. As a result, $G_{\mathcal{V}}$ contains an equal or larger number of branching, i.e. unitig-flanking vertices

[1] We verified this behavior of BCALM 2 through communicating the authors.
[2] Maximal unitigs with at least one ending side having no edge.
[3] For clarity, we are not considering the $k$-mer orientations.

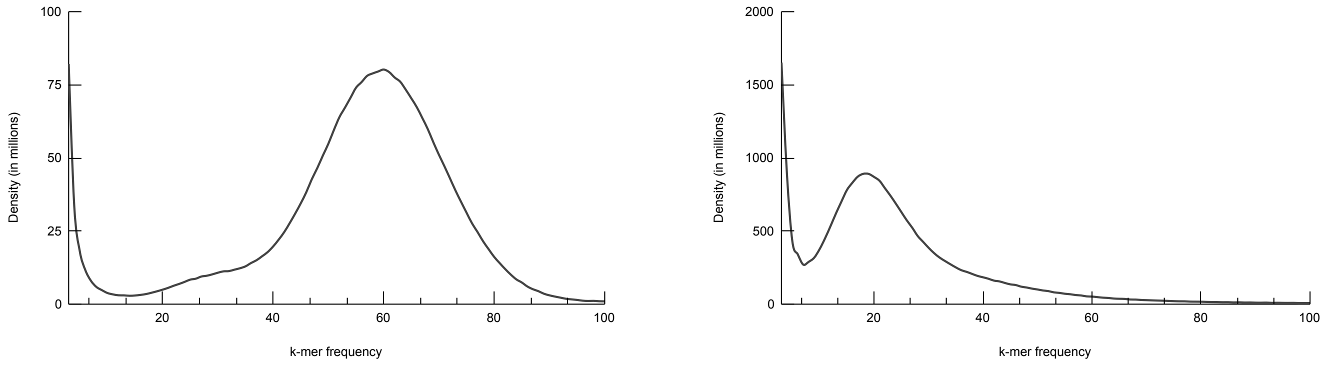


**(a)** **(b)**

**Figure 1:** Frequency distribution of k-mer abundances: (a) for the human read set NIST HG004 with $k = 27$, and (b) for the white spruce read set NCBI PRJNA83435 with $k = 55$. Densities for the frequencies 1 and 2 have been omitted from the plots, as those dwarf the other frequency densities and skew the plots drastically.

**Table 1:** Time-, memory-, and disk-performance results for constructing compacted de Bruijn graphs from short-read sets.

| Dataset | k | Thread-count | ABySS-Bloom-dBG: Small-memory | ABySS-Bloom-dBG: Large-memory | Bifrost | deGSM | BCALM 2 | Cuttlefish 2: Default memory | Cuttlefish 2: Match second-best memory | Cuttlefish 2: Un-restricted memory |
|---|---|---|---|---|---|---|---|---|---|---|
| Human | 27 | 8 | 22h 18m (39.3 \| 0) | 20h 23m (71.3 \| 0) | 11h 43m (48.5 \| 0) | 10h 36m (235.8 \| 737) | 04h 23m (6.7 \| 344) | **01h 13m** (**3.2** \| 209) | 01h 10m (6.2 \| 209) | 01h (11.3 \| 186) |
| | | 16 | 11h 38m (39.3 \| 0) | 11h 02m (71.3 \| 0) | 09h 39m (48.6 \| 0) | 07h 08m (235.8 \| 730) | 04h 58m (8.9 \| 342) | **56m** (**3.3** \| 209) | 56m (7.6 \| 209) | 51m (11.3 \| 186) |
| | 55 | 8 | 16h 32m (34.0 \| 0) | 15h 58m (66.0 \| 0) | 05h 43m (43.8 \| 0) | 16h 50m (293.2 \| 1147) | 04h 01m (7.4 \| 296) | **02h 20m** (**3.5** \| 147) | 01h 08m (7.1 \| 147) | 01h 03m (11.3 \| 142) |
| | | 16 | 09h 28m (34.1 \| 0) | 08h 37m (66.1 \| 0) | 04h 16m (43.9 \| 0) | 15h 54m (293.3 \| 1147) | 04h 26m (10.5 \| 293) | **02h 02m** (**3.7** \| 147) | 01h 11m (9.5 \| 147) | 51m (11.3 \| 142) |
| Human RNA-seq | 27 | 8 | 11h 47m (33.7 \| 0) | 11h 22m (65.7 \| 0) | 06h 04m (7.2 \| 0) | 01h 35m (87.1 \| 31) | 02h 58m (3.8 \| 217) | **30m** (**2.9** \| 89) | – | 18m (80.1 \| 85) |
| | | 16 | 11h 38m (39.3 \| 0) | 07h 38m (65.7 \| 0) | 07h 24m (7.2 \| 0) | 01h 37m (87.2 \| 31) | 02h 46m (3.9 \| 217) | **20m** (**3.0** \| 89) | – | 12m (80.1 \| 85) |
| Gut microbiome | 27 | 16 | 18h 47m (42.0 \| 0) | 20h 12m (74.0 \| 0) | 03h 54m (38.1 \| 0) | 02h 28m (157.2 \| 362) | 02h 34m (7.7 \| 165) | **26m** (**3.5** \| 78) | 23m (6.7 \| 78) | 20m (26.8 \| 69) |
| | 55 | | 1d 17h 43m (35.9 \| 0) | 1d 08h 09m (67.8 \| 0) | 02h 44m (46.7 \| 0) | 06h 53m (293.3 \| 624) | 03h 02m (12.5 \| 158) | **44m** (**4.0** \| 52) | 25m (11.3 \| 52) | 20m (69.9 \| 50) |
| Soil | 27 | 16 | 1d 18h 35m (150.4 \| 0) | 14h 24m (275.0 \| 0) | 15h 28m (274.1 \| 0) | 1d 14h 29m (235.8 \| 3287) | 19h 39m (52.0 \| 681) | **02h 01m** (**19.2** \| 161) | 02h 18m (40.9 \| 161) | 01h 35m (40.9 \| 210) |
| | 55 | | 07h 57m (128.9 \| 0) | 06h 36m (256.8 \| 0) | 05h 49m (157.0 \| 0) | 1d 11h 05m (293.3 \| 2959) | 08h 30m (27.5 \| 419) | **03h 02m** (**11.1** \| 132) | 02h 43m (23.3 \| 132) | 01h 38m (23.3 \| 234) |
| White spruce | 27 | 16 | * | X | X | † | 2d 06h 12m (36.8 \| 2171) | **10h 05m** (**14.0** \| 1362) | 07h 47m (35.2 \| 1362) | 07h 13m (204.2 \| 1208) |
| | 55 | | * | X | X | † | 2d 09h 59m (31.6 \| 1505) | **10h 12m** (**23.8** \| 897) | 10h 08m (31.1 \| 897) | 07h 24m (279.3 \| 865) |

Each cell contains the running time in wall clock format, and in parentheses: the maximum memory usage and the maximum intermediate disk-usage separated by |, in gigabytes. All the execution details and other relevant information can be found in Table 1 (see main text).

**Table 2:** Time- and memory-performance results for constructing compacted de Bruijn graphs from the human read set NIST HG004, with frequency threshold $f_0 = 2$.

| k | Thread-count | ABySS-Bloom-dBG: Small-memory | ABySS-Bloom-dBG: Large-memory | Bifrost | deGSM | BCALM 2 | Cuttlefish 2: Default memory | Cuttlefish 2: Match second-best memory | Cuttlefish 2: Un-restricted memory |
|---|---|---|---|---|---|---|---|---|---|
| 27 | 8 | Δ | 1d 16h 16m (77.5) | 11h 43m (48.5) | 09h 34m (235.8) | 06h 01m (8.9) | **01h 15m** (**3.9**) | 01h 08m (8.6) | 01h 02m (11.3) |
| | 16 | 1d 14h 08m (46.9) | 1d 02h 10m (77.5) | 11h 02m (48.6) | 08h 24m (235.8) | 06h 19m (11.6) | **57m** (**4.1**) | 52m (11.4) | 49m (11.3) |
| 55 | 8 | Δ | 1d 08h 20m (67.1) | 05h 43m (43.8) | 17h 23m (293.2) | 05h 51m (7.7) | **02h 21m** (**4.1**) | 01h 10m (8.5) | 01h 04m (11.3) |
| | 16 | Δ | 16h 29m (67.1) | 04h 16m (43.9) | 15h 31m (293.2) | 06h 08m (10.6) | **02h 05m** (**4.3**) | 01h (10.4) | 46m (11.3) |

Each cell contains the running time in wall clock format, and the maximum memory usage in gigabytes, in parentheses. Details on executing the different tool implementations can be found in Table 1 (See main text).
The best performance with respect to each metric in each row is highlighted, where only the default-memory mode is considered for Cuttlefish 2. The Δ's in the ABySS-Bloom-dBG results denote that the corresponding executions were allowed to run for at least 2 days, before being explicitly terminated.

**Table 3:** Time-, memory-, disk-performance results for constructing compacted de Bruijn graphs from whole-genome reference collections.

| Dataset (genome count) | $k$ | Thread-count | Bifrost | deGSM | BCALM 2 | Cuttlefish 2 | |
|---|---|---|---|---|---|---|---|
| | | | | | | Default memory | Unrestricted memory |
| Human gut (30K) | 27 | 8 | 06h (155.1 \| 0) | Δ | 10h 06m (21.5 \| 473) | **01h 39m** (**15.2** \| 111) | 01h 39m (32.5 \| 183) |
| | | 16 | 05h 30m (155.1 \| 0) | | 09h 05m (22.0 \| 473) | **01h 01m** (**15.5** \| 111) | 59m (32.5 \| 183) |
| | 55 | 8 | 08h 47m (279.2 \| 0) | | 11h 49m (**18.6** \| 708) | **04h 14m** (20.6 \| 262) | 03h 42m (44.4 \| 480) |
| | | 16 | 08h 20m (279.2 \| 0) | | 09h 45m (**19.2** \| 708) | **03h 50m** (20.9 \| 262) | 03h 10m (44.3 \| 480) |
| Human (100) | 27 | 8 | 35h 45m (355.9 \| 0) | 19h 23m (235.8 \| 1219) | ‡ | **04h 32m** (**27.7** \| 311) | 04h 09m (59.7 \| 345) |
| | | 16 | 32h 14m (355.9 \| 0) | 14h 07m (235.8 \| 1260) | ‡ | **03h 19m** (**28.1** \| 311) | 02h 49m (59.7 \| 345) |
| | 55 | 8 | * | † | 2d 23h 31m (302.9 \| 2150) | **15h 08m** (**56.0** \| 1288) | 13h 47m (121.8 \| 1332) |
| | | 16 | * | † | * | **12h** (**56.2** \| 1288) | 11h 33m (121.8 \| 1332) |
| Bacterial archive (661K) | 27 | 16 | X | X | ‡ | **16h 38m** (**48.7** \| 2658) | 16h 24m (104.9 \| 2347) |
| | 55 | | | | 4d 10h 11m (63.3 \| 2212) | **22h 44m** (**59.9** \| 2047) | 22h 20m (129.5 \| 1974) |

Each cell contains the running time in wall clock format, and in parentheses: the maximum memory usage and the maximum intermediate disk-usage separated by |, in gigabytes. All the execution details and other relevant information can be found in Table 2 (see main text).

**Table 4:** Timing performance for constructing compacted de Bruijn graphs, excluding the initial $k$-mer (or $(k+1)$-mer) enumeration step.

| Dataset | $k$ | Thread-count | BCALM 2 | Cuttlefish 2 | |
|---|---|---|---|---|---|
| | | | | Default memory | Unrestricted memory |
| Short-read sets | | | | | |
| Human | 27 | 8 | 01h 09m | 13m | 11m |
| | | 16 | 01h 06m | 09m | 07m |
| | 55 | 8 | 01h 12m | 23m | 17m |
| | | 16 | 01h 14m | 15m | 14m |
| Human RNA-seq | 27 | 8 | 23m | 02m | 01m |
| | | 16 | 22m | 01m | 01m |
| Gut microbiome | 27 | 16 | 01h 38m | 09m | 07m |
| | 55 | | 01h 46m | 16m | 13m |
| Soil | 27 | 16 | 04h 56m | 01h 17m | 01h 04m |
| | 55 | | 05h 09m | 02h 17m | 01h 11m |
| White spruce | 27 | 16 | 18h 24m | 01h 30m | 52m |
| | 55 | | 17h 17m | 04h 47m | 03h 15m |
| Whole-genome reference collections | | | | | |
| Human gut (30K) | 27 | 8 | 08h 24m | 01h 22m | 01h 24m |
| | | 16 | 07h 25m | 49m | 46m |
| | 55 | 8 | 09h 09m | 03h 44m | 03h 15m |
| | | 16 | 07h 34m | 03h 50m | 02h 45m |
| Human (100) | 27 | 8 | ‡ | 03h | 02h 41m |
| | | 16 | ‡ | 02h 22m | 01h 31m |
| | 55 | 8 | 2d 06h 50m | 13h 17m | 11h 50m |
| | | 16 | * | 10h 16m | 09h 25m |
| Bacterial archive (661K) | 27 | 16 | ‡ | 04h 16m | 03h 26m |
| | 55 | | 1d 08h 36m | 10h 50m | 10h 32m |

Each cell contains the running time in wall clock format, excluding the times incurred by the initial: (a) $k$-mer enumeration step of BCALM 2, and (b) $(k+1)$-mer enumeration step of Cuttlefish 2. All the execution details and other relevant information can be found in the Tables 1 and 2 (see main text).

than $G_{\mathcal{E}}$. [4] This reduces the number of maximal unitigs reported in in edge-centric de Bruijn graphs compared to vertex-centric ones.

***Vertex-filtering versus Edge-filtering.*** Related to the above point, since the fundamental units of de Bruijn graph construction in Cuttlefish 2 are the edges, this is where error-filtering is performed prior to construction. Conversely, BCALM 2 and Bifrost take a vertex-centric approach to construction and hence filtering is performed on the vertex set. In some corner cases, where unit-abundances are very close to the selected threshold, this can lead to small-scale differences in which $k$-mers are filtered out prior to construction.

Consider a given threshold $f_0$. Any $k$-mer $x$ present in the input at least $f_0$ times is a vertex in the vertex-centric graph. But there may not exist any $(k+1)$-mer $z$ in the input that occurs at least $f_0$ times and contains $x$, thus $x$ is absent as a vertex in the corresponding edge-centric graph. On the opposite direction, any $k$-mer $y$, substring of a $(k+1)$-mer $z$ that is present in the input at least $f_0$ times, is a vertex in the

[4] It could be possible that some of these additional edges in $G_{\mathcal{V}}$ connect two separate maximal unitigs into one, thus actually reducing branching vertices. The assumption that $G_{\mathcal{E}}$ contains no tips prevents this—there can not exist an edge $\{x, y\}$ in $G_{\mathcal{V}}$ such that, $x$ and $y$ are tip-ends in $G_{\mathcal{E}}$, and connect the two tips they belong to into one single maximal unitig in $G_{\mathcal{V}}$.

edge-centric graph. It also implies that $y$ has an abundance of at least $f_0$ in the input, and thus is also a vertex in the vertex-centric graph.
Therefore, when using a same frequency threshold $f_0$ for k-mers and $(k+1)$-mers, vertex-centric de Bruijn graphs must always have an equal or greater number of vertices than edge-centric ones.

**1.6. Compacted de Bruijn graph properties.** Suppl. Table 5 contains some notable characteristics of the original de Bruijn graphs and their compacted forms.

**1.7. Maximal path cover construction.** Suppl. Table 6 provides a comparison of the maximal unitig based and the maximal path cover based representations of the de Bruijn graphs.

**1.8. Parallel scaling.** Suppl. Fig. 2 demonstrates the timing-profile and speedup for each step of CUTTLEFISH 2, on the same setting as described in Sec. 2.7 (see main text), but with $k = 55$.

**1.9. Application in associative $k$-mer index construction.** The utility of CUTTLEFISH 2 and any compacted de Bruijn graph constructor depends upon the downstream applications for which it is used. In this proof of-concept section, we demonstrate the improvement provided by CUTTLEFISH 2 over alternative methods in a pipeline that constructs an associative k-mer index over a collection of reads or references. These indices, sometimes implicitly, form a fundamental component in various computational genomics tasks, such as in tools for variant detection and genotyping (4), RNA isoform quantification (5), large-scale sequence search (6), and k-mer abundance indexing (7).
Given a set $\mathcal{V}$ of k-mers, an associative k-mer index of $\mathcal{V}$ consists of a bijective mapping $f : \mathcal{V} \to [0, |\mathcal{V}|)$. It is different from a minimal perfect hash $h : \mathcal{V} \to [0, |\mathcal{V}|)$ in that, any alien k-mer $v \notin \mathcal{V}$ can be detected by $f$ as absent in $\mathcal{V}$, i.e. $\forall v \notin \mathcal{V}\ f(v) = -1$; but not necessarily by $h$.
We investigated the overall performance difference for a pipeline that uses SSHASH (8) to index the k-mer set—represented with the de Bruijn graph—of several datasets of various sizes. In the pipeline, we first extract the maximal path (or unitig) sequences from the graph, and then use SSHASH to construct a k-mer index from these sequences. We compare the performance of this pipeline when using CUTTLEFISH 2 to extract the maximal path cover (or unitigs) versus using UST (or BCALM 2). Suppl. Table 7 provides a comparison of the performances.
We observe that, for intermediate or large datasets, when using UST (or BCALM 2) to extract the maximal path cover (or unitigs), the extraction of the sequences itself is both the time and the memory bottleneck. That is, indexing the extracted sequences with SSHASH is both faster and more memory frugal than extracting the sequences in the first place, often by a considerable factor. On the other hand, when we use CUTTLEFISH 2 to extract the sequences, the time taken to extract the sequences, and therefore the time taken to construct the entire index, is reduced dramatically. Additionally, the memory usage of CUTTLEFISH 2 is often comparable (or in the case of the Gut microbiome reads less than) that of SSHASH. Thus, replacing UST (or BCALM 2) with CUTTLEFISH 2 in this index construction greatly reduces the bottleneck step of associative index construction, and sometimes even shifts the bottleneck itself from the task of extracting a maximal path cover (or unitigs) of the de Bruijn graph to the task of constructing the index from these sequences.

**1.10. Tools and execution commands.** For the experiments, we used the following versions of the tools—(1) ABYSS-BLOOM-DBG from ABYSS 2.0 (v2.3.1) (2) BCALM 2 (v2.2.3) (3) BIFROST (v1.0.6.5) (4) DEGSM (v1.0) (5) PROPHASM (v0.1.1) (6) UST from ESS-COMPRESS (v2.1), (7) SSHASH (v2.1.0), and (8) CUTTLEFISH 2 (commit ID 0a049a5).
The following commands have been used in executing the tools.

- ABYSS-BLOOM-DBG:
```
abyss-bloom-dbg -b${bf_size}
-H${bf_hash_num} -j${threads}
-k${k} -kc=${min_count}
-out=${op_file} -v ${ip_files}
```
- BCALM 2:
```
bcalm -in ${ip_list}
-kmer-size ${k}
-abundance-min ${min_count}
-nb-cores ${threads}
-max-memory ${memory}
-max-disk ${disk}
-out-tmp ${temp_dir}
-out ${op_file}
```
- BIFROST:
```
Bifrost build
-${ip_type_arg} ${ip_list} -k ${k}
-t ${threads} -o ${op_file} -v
```
where `ip_type_arg` is either `r` or `s`, based on whether reference-sequences or short-reads are provided as input, respectively.
- DEGSM:
```
LD_LIBRARY_PATH=${jellyfish_lib_path}
deGSM -k ${k} ${min_count_arg}
-t ${threads} -m ${memory}G
${zipped_arg} ${jellyfish_lib_path}
${op_file}.bwt ${ip_dir}
```
and
```
ubwt unipath ${op_file}.bwt
-t ${threads} -o ${op_file}
```
where `min_count_arg` is `-l ${min_count}` or empty, based on whether short-reads or reference-sequences are provided as input, respectively; and `${zipped_arg}` is `-g` if the input files are in `.gz` format, and empty otherwise.

**Table 5:** Some properties of the ordinary de Bruijn graph and its compacted form.

| Dataset | k | de Bruijn graph # Vertices ($\times 10^6$) | # Edges ($\times 10^6$) | Compacted de Bruijn graph # Vertices ($\times 10^6$) | # Edges ($\times 10^6$) | Max. unitig length (kbp) |
|---|---|---|---|---|---|---|
| | | **Short-read sets** | | | | |
| Human | 27 | 2,490 | 2,514 | 58 | 81 | 21.0 |
| | 55 | 2,867 | 2,874 | 24 | 31 | 36.7 |
| Human RNA-seq | 27 | 320 | 309 | 66 | 55 | 1.1 |
| Gut microbiome | 27 | 2,580 | 2,616 | 205 | 241 | 5.6 |
| | 55 | 3,107 | 3,111 | 167 | 172 | 3.9 |
| Soil | 27 | 16,522 | 16,335 | 939 | 752 | 3.1 |
| | 55 | 9,391 | 9,121 | 432 | 162 | 2.1 |
| White spruce | 27 | 11,236 | 11,694 | 1,244 | 1,702 | 7.6 |
| | 55 | 20,536 | 20,725 | 704 | 894 | 17.0 |
| | | **Whole-genome references** | | | | |
| Human gut (30K) | 27 | 13,132 | 13,340 | 569 | 776 | 92.7 |
| | 55 | 17,901 | 18,048 | 437 | 584 | 368.4 |
| Human (100) | 27 | 24,094 | 25,055 | 2,315 | 3,276 | 8.0 |
| | 55 | 49,220 | 50,039 | 2,122 | 2,941 | 19.2 |
| Bacterial archive (661K) | 27 | 42,330 | 42,871 | 1,437 | 1,978 | 114.2 |
| | 55 | 52,288 | 52,542 | 749 | 1,003 | 679.6 |

**Table 6:** Comparison of the maximal unitig based and the maximal path-cover based representations of de Bruijn graphs.

| Dataset | k | k-mer-count | Maximal Unitigs # Unitigs | Avg. length (bp) | Max. length (bp) | base/k-mer | Maximal Path-cover # Paths | Avg. length (bp) | Max. length (bp) | base/k-mer |
|---|---|---|---|---|---|---|---|---|---|---|
| | | | **Short-read sets** | | | | | | | |
| Roundworm | 27 | 93,574,387 | 608,793 | 179.7 | 46,859 | 1.17 | 218,508 | 454.2 | 63,884 | 1.06 |
| | 55 | 96,582,016 | 292,444 | 384.3 | 66,206 | 1.16 | 129,203 | 801.5 | 79,500 | 1.07 |
| Gut microbiome | 27 | 2,579,749,776 | 204,893,577 | 38.6 | 5,633 | 3.07 | 97,631,499 | 52.4 | 6,871 | 1.98 |
| | 55 | 3,106,506,224 | 167,337,716 | 72.6 | 3,857 | 3.91 | 91,760,241 | 87.9 | 6,058 | 2.60 |
| Human | 27 | 2,490,358,687 | 57,804,370 | 69.1 | 21,012 | 1.60 | 19,811,145 | 151.7 | 21,066 | 1.21 |
| | 55 | 2,866,610,943 | 23,778,178 | 174.6 | 36,697 | 1.45 | 8,915,957 | 375.5 | 48,560 | 1.17 |
| | | | **Whole-genome references** | | | | | | | |
| Roundworm | 27 | 93,471,568 | 527,960 | 203 | 75,221 | 1.15 | 173,552 | 564.6 | 78,941 | 1.05 |
| | 55 | 96,417,950 | 165,081 | 638.1 | 130,760 | 1.16 | 55,385 | 1,794.9 | 130,767 | 1.03 |
| Human | 27 | 2,431,778,046 | 44,459,296 | 80.7 | 29,022 | 1.48 | 14,209,926 | 197.1 | 29,034 | 1.15 |
| | 55 | 2,737,097,058 | 12,522,233 | 272.6 | 94,673 | 1.25 | 4,071,450 | 726.3 | 123,699 | 1.08 |
| 7 humans | 27 | 2,498,416,058 | 54,440,059 | 71.9 | 18,424 | 1.57 | 17,507,551 | 168.7 | 30,285 | 1.18 |
| | 55 | 2,907,442,632 | 23,169,472 | 179.5 | 33,969 | 1.43 | 7,608,240 | 436.1 | 44,620 | 1.14 |

Given a de Bruijn graph $G(\mathcal{R}, k) = (\mathcal{V}, E)$ and a representation of it $\mathcal{P}$, the *base/k-mer* metric is computed as $\sum_{p \in \mathcal{P}} |p| \,/\, |\mathcal{V}|$, i.e. the number of nucleobase characters required in average per k-mer for the literal representation of the paths in $\mathcal{P}$ (maximal unitigs decomposition is also a path-cover). If the 2-bit/nucleobase encoding is used instead of the literal representations, then then the *bits/k-mer* requirement would be 1/4'th of the base/k-mer requirement.

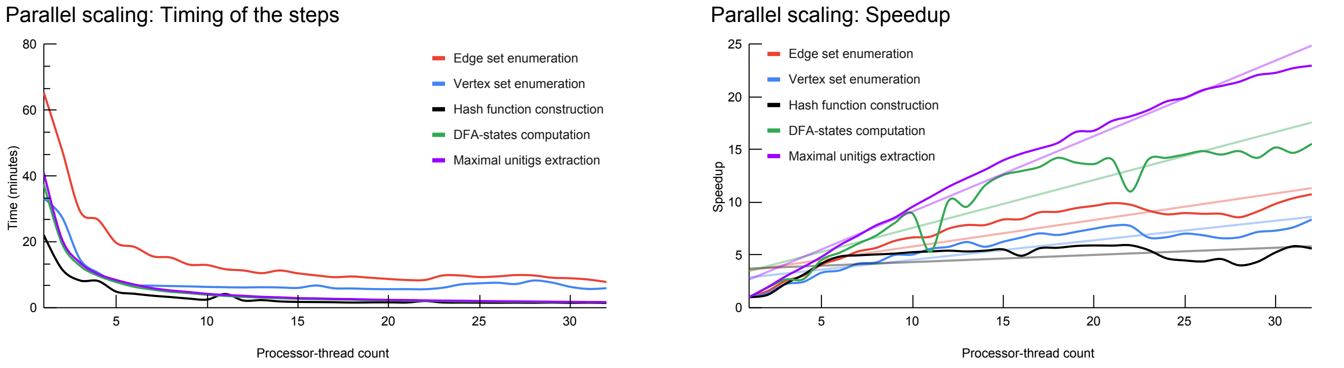


**(a)** Time incurred by each individual step. **(b)** Speedup of each individual step.

**Figure 2:** Parallel-scaling metrics for CUTTLEFISH 2 across 1–32 processor threads, using $k = 55$ on the (downsampled) human read set NIST HG004, with the frequency threshold $f_0 = 3$.

- PROPHASM:
  `prophasm -k ${k} ${io_paths}`
  where `${io_paths}` is a concatenation of the following: `-i ${file_name} -o ${op_file}`, repeated for each different input file, with the same output file.

- UST:

**Table 7:** Time- and memory-performance results for constructing an associative k-mer index using the de Bruijn graph.

| Dataset | # k-mers ($\times 10^6$) | Maximal path cover construction | | Index construction | Full pipeline | |
|---|---|---|---|---|---|---|
| | | UST | CUTTLEFISH 2 | SSHASH | UST-total | CUTTLEFISH 2-total |
| Human reads | 2,490 | 04h 56m (13.1) | 01h 18m (3.2) | 35m (3.0) | 05h 31m (13.1) | 01h 53m (3.2) |
| Gut micro-biome reads | 2,580 | 03h 10m (39.2) | 53m (3.3) | 26m (5.9) | 03h 36m (39.2) | 01h 20m (5.9) |
| Human genome ref. | 2,432 | 01h 09m (10.3) | 14m (3.2) | 25m (3.0) | 01h 34m (10.3) | 39m (3.2) |
| 7 human refs. | 2,498 | 01h 49m (20.2) | 18m (3.2) | 25m (3.0) | 02h 15m (20.2) | 43m (3.2) |
| | | Maximal unitigs construction | | Index construction | Full pipeline | |
| | | BCALM 2 | CUTTLEFISH 2 | SSHASH | BCALM 2-total | CUTTLEFISH 2-total |
| Human reads | 2,490 | 04h 58m (8.9) | 56m (3.3) | 27m (4.0) | 05h 25m (8.9) | 01h 23m (4.0) |
| Human RNA-seq | 320 | 02h 46m (3.9) | 20m (3.0) | 04m (2.0) | 02h 50m (3.9) | 24m (3.0) |
| Gut micro-biome reads | 2,580 | 02h 34m (7.7) | 26m (3.5) | 33m (7.9) | 03h 07m (7.9) | 59m (7.9) |

Each cell contains the running time in wall clock format, and the maximum memory usage in gigabytes, in parentheses. In all the executions, k is 27, and the maximal path cover and the maximal unitigs are the ones obtained from the experiments performed in Secs. 2.3, 2.4, and 2.5 (see main text). The dataset descriptions are also present in these sections. All the execution details and other relevant information can be found in Tables 1, 2, and 3 (see main text). The thread count used for UST and CUTTLEFISH 2 in the maximal path cover based construction is 8; for BCALM 2 and CUTTLEFISH 2 in the maximal unitigs based construction, it is 16. The SSHASH implementation is single-threaded.

```
essCompress -i ${ip_list} -k ${k}
-a ${min_count} -t ${threads}
-o ${op_dir} -u -v
```
where the `-t ${threads}` argument has been added by us to control the number of processor-cores for it to use—its default setting uses up-to all the available cores.

- SSHASH:
```
cgexec -g memory:${cgroup_name}
sshash/build/build ${input} ${k}
${minimizer_len} -o ${output}
-d ${temp_dir} --verbose
```
where the `cgroup_name` is a Linux control group, set with an appropriate memory limit to restrict SSHASH's reported memory usage. This is necessary due to SSHASH's use of the `mmap` system call, which results in the counting of unused shared pages toward the program's memory usage, particularly in a memory-rich execution environment where the program is not under memory pressure. As such, the `time` command will report a much higher memory usage than is actually required by SSHASH to run. The `cgroup` execution places a hard limit on the memory the program can use, and applies the requisite memory pressure to ensure that the reported memory usage is much closer to what is actually required for successful execution.

- CUTTLEFISH 2:
In the following, the `${read_or_ref_arg}` is either `read` or `ref`, based on whether reference-sequences or short-reads are provided as input, respectively.
  - Compacted de Bruijn graph construction (with default memory):
```
cuttlefish build
--${read_or_ref_arg}
-l ${ip_list} -k ${k}
-c ${min_count} -t ${threads}
-w ${temp_dir} -o ${op_prefix}
```
  - Compacted de Bruijn graph construction (with a given memory threshold):
```
cuttlefish build
--${read_or_ref_arg}
-l ${ip_list} -k ${k}
-c ${min_count} -t ${threads}
-m ${memory}
-w ${temp_dir} -o ${op_prefix}
```
  - Compacted de Bruijn graph construction (with unrestricted memory):
```
cuttlefish build
--${read_or_ref_arg}
-l ${ip_list} -k ${k}
-c ${min_count} -t ${threads}
--unrestrict-memory
-w ${temp_dir} -o ${op_prefix}
```
  - Maximal path-cover construction (with default memory):
```
cuttlefish build
--${read_or_ref_arg}
-l ${ip_list} -k ${k}
-c ${min_count} -t ${threads}
-w ${temp_dir} -o ${op_prefix}
--path-cover
```
  - Maximal path-cover construction (with unrestricted memory):

```
cuttlefish build
--${read_or_ref_arg}
-l ${ip_list} -k ${k}
-c ${min_count} -t ${threads}
--unrestrict-memory
-w ${temp_dir} -o ${op_prefix}
--path-cover
```

The scripts used to perform the experiments described in the paper are available at https://github.com/COMBINE-lab/cuttlefish_experiments.

### 2. Methods

**2.1. Upgrades in the KMC 3 algorithm.** We implemented several upgrades in the KMC 3 algorithm to tune it to the efficiency needs for CUTTLEFISH 2. Here we discuss those briefly. Although the upgrades were designed specifically for usage in CUTTLEFISH 2, those may also be suitable in other bioinformatics pipelines, and are publicly available in the KMC 3 GitHub repository (https://github.com/refresh-bio/kmc).

***2.1.1. Counting $k$-mers from existing KMC 3 database.*** KMC 3 is updated so as to be able to count $k'$-mers from a $k$-mer database produced by another KMC 3 execution, for some $k' < k$. This allows reducing computational resources needed to determine the set of vertices, $\mathcal{V}$, as it may be directly computed from the set of edges, $\mathcal{E}$, without the need of an entire pass over all the input sequences. This is especially relevant in the case of sequencing reads. Technically, the KMC 3 API is used in the listing mode to enumerate all $k$-mers that are further processed as if they were reads.

***2.1.2. Estimate $k$-mer abundance histogram during the first stage in KMC 3.*** This upgrade allows efficient estimation of the total number of unique $k$-mers present in the input during the first stage of KMC 3. The estimation is performed by our optimized implementation of the NTCARD algorithm (1).

***2.1.3. Using KMC 3 directly from C++ code with API.*** A new API to use KMC 3 directly from inside some C++ code is designed for CUTTLEFISH 2, and it is usable in general. Furthermore, it is possible to set parameters for the second stage of KMC 3 based on the results of the first stage. The detailed documentation of API is available in the KMC 3 GitHub repository: https://github.com/refresh-bio/kmc/wiki/Use-the-KMC-directly-from-code-through-the-API. Combining this with the capability to estimate $k$-mer abundance histograms, it is possible to bound the memory-usage of the second stage of KMC 3 such that it uses at most the peak amount of memory required in the next steps of CUTTLEFISH 2.

***2.1.4. Storing $k$-mers without counts in KMC 3 databases.*** This upgrade affects disk usage. To date, KMC 3 output required at least one byte per $k$-mer to store a counter. In some applications, e.g. to build the compacted de Bruijn graph without abundance estimates for the vertices, the counters are not required and can be skipped. In practice, this leads to the reduction of disk usage and, as a consequence, reduction in the total I/O costs, which in turn affects the running time.

### 3. Proofs

***Lemma** 1:* The $(k+1)$-mers $z$ and $\overline{z}$ induce the same bidirected edge in a de Bruijn graph $G(\mathcal{S}, k)$.

*Proof:* Consider a $(k+1)$-mer $z$ from some input string $s \in \mathcal{S}$. Let $x = \mathrm{pre}_k(z)$ and $y = \mathrm{suf}_k(z)$. Then $z$ can be expressed as $z = x \odot^{k-1} y$.

$z$ induces an edge between the vertices $\widehat{x}$ and $\widehat{y}$. It is incident to the $\mathtt{back}$ of $\widehat{x}$ when $\widehat{x} = x$ holds, and is incident to the $\mathtt{front}$ when $\widehat{x} = \overline{x}$ (see Sec. 3.2, main text).

$z$'s reverse complement is $\overline{z} = \overline{y} \odot^{k-1} \overline{x}$, and it induces an edge between $\widehat{y}$ and $\widehat{x}$. It is incident to the $\mathtt{front}$ of $\widehat{x}$ if $\widehat{x} = \overline{x}$ holds, and is incident to the $\mathtt{back}$ if $\widehat{x} = x$—the same side as that of $z$'s edge.

It can be proven likewise that the edges are incident to the same side of $\widehat{y}$. Therefore, $z$ and $\overline{z}$ induce edges between the same vertex-pair $\{\widehat{x}, \widehat{y}\}$, incident to the same sides—inducing the same bidirected edge. ■

***Lemma** 2:* A side of a vertex can have at most $|\Sigma|$ distinct edges in a de Bruijn graph $G(\mathcal{S}, k)$.

*Proof:* Consider a vertex $\widehat{v}$ in $G(\mathcal{S}, k)$. WLOG, we prove the claim for the $\mathtt{back}$ of $\widehat{v}$.

An edge $e$ connected to $\widehat{v}$ and induced by a $(k+1)$-mer $z$ is incident to $\widehat{v}$'s $\mathtt{back}$ iff: (1) $\mathrm{pre}_k(z) = \widehat{v}$; or (2) $\mathrm{suf}_k(z) = \overline{\widehat{v}}$ (see Sec. 3.2, main text). For case (1), the possible $z$'s form the set $\mathcal{E}_1 = \{\widehat{v} \cdot c : c \in \Sigma\}$. For case (2), the set is $\mathcal{E}_2 = \{c' \cdot \overline{\widehat{v}} : c' \in \Sigma\}$, which is the same as $\{\overline{c} \cdot \overline{\widehat{v}} : c \in \Sigma\}$, letting $c' = \overline{c}$ [5].

As per Lemma 1, the $(k+1)$-mers $c \cdot \widehat{v}$ and $\overline{\widehat{v}} \cdot \overline{c}$ induce the same bidirected edge, where $c \in \Sigma$. Thus $\mathcal{E}_1$ and $\mathcal{E}_2$ induce the same set of edges. Therefore, the $\mathtt{back}$ of $\widehat{v}$ can have at most $|\mathcal{E}_1| = |\mathcal{E}_2| = |\Sigma|$ distinct edges. ■

***Lemma** 3:* A vertex $\widehat{v}$ is noted to be a flanking vertex in a de Bruijn graph $G(\mathcal{S}, k)$ iff it is an endpoint of a maximal unitig.

*Proof:* Let $\mathcal{C}_v$ be the state-class of $\widehat{v}$'s automaton and $p$ be the maximal unitig containing $\widehat{v}$. The term *branching* in the proof means connecting to multiple distinct edges.

First, assume that $\widehat{v}$ is marked as a flanking vertex. We prove that $\widehat{v}$ is an endpoint of $p$. As per the definition of flanking vertices (see Sec. 3.3.8, main text), either of the following holds:

1. $\mathcal{C}_v$ is not *unique-front unique-back*. Then from Corollary 1, $\widehat{v}$ has at least one side $s_v$ with either $0$ or $> 1$ distinct edges. It is not possible to extend $p$ through $s_v$—either there is no edge, or the addition introduces an internal branching vertex $\widehat{v}$ in $p$.

2. $\mathcal{C}_v$ is *unique-front unique-back*, and a side of it, $s_v$, is connected to a branching side $s_u$ of a vertex $\widehat{u}$. Then $p$ can not be extended through $s_v$, because the extension includes $s_u$ as an internal side to $p$, which is branching.

[5] As per our definitions, the set $\Sigma$ of symbols is closed under complementing.

In either case, $\hat{v}$ is an endpoint of $p$.

Now assume that $\hat{v}$ is an endpoint of $p$. We prove that $\hat{v}$ is marked as a flanking vertex. Based on the adjacencies of $\hat{v}$, either of the following holds:

1. $\hat{v}$ has at least one side $s_v$ that is either empty or branching. From Corollary 1, $C_v$ is not *unique-front unique-back*.

2. $\hat{v}$ has one unique edge at each side. Say that its side $s_v$ restricts $p$ from extending farther, and $s_v$ connects to the side $s_u$ of a vertex $\hat{u}$. The definition of unitigs implies that $s_u$ must be branching. This in turn implies from Corollary 1 that $\hat{u}$'s automaton's state is from the state-class: (i) either *fuzzy-front fuzzy-back*; or (ii) *fuzzy-front unique-back*, in which case $s_u$ is $\mathtt{front}$; or (iii) *unique-front fuzzy-back*, in which case $s_u$ is $\mathtt{back}$.

In either case, $\hat{v}$ fulfills the conditions for being a flanking vertex. ■

*Corollary 1:* For the automaton $M_v$ of a vertex $\hat{v}$ in a de Bruijn graph $G(\mathcal{S}, k)$, applying $\delta$ on $M_v$ with all the incident edges of $\hat{v}$ (in any order) transitions its state from $q_0$ to $q_v$ belonging to the state-class $C_v$, such that $C_v$ is:

1. *fuzzy-front fuzzy-back*, iff $\hat{v}$ does not have exactly one unique edge at any of its sides

2. *fuzzy-front unique-back*, iff $\hat{v}$ has exactly one unique edge only at its $\mathtt{back}$

3. *unique-front fuzzy-back*, iff $\hat{v}$ has exactly one unique edge only at its $\mathtt{front}$

4. *unique-front unique-back*, iff $\hat{v}$ has exactly one unique edge at each of its sides.

*Proof:* The proof is trivial from the definition of the transition function $\delta$, illustrated in detail in Fig. 4 (see main text). ■

*Theorem 1:* CUTTLEFISH $2(\mathcal{R}, k, f_0)$ is correct.

*Proof:* Following from Corollary 1, the COMPUTE-AUTOMATON-STATES algorithm correctly computes the state-classes of all the automata. Besides, CUTTLEFISH 2's modeling scheme of a vertex $\hat{v}$ with an automaton $M_v$ ensures that if a side $s_v$ has a unique incident edge $e$, an encoding of $e$ is preserved in $M_v$'s state $q_v$, observable from the illustration of $\delta$ in Fig. 4 (see main text). Hence, all the internal edges of the maximal unitigs are retained within the states.

For some vertex $\hat{v} \in \mathcal{V}$, let $p$ be the maximal unitig containing $\hat{v}$, and $p = (\hat{v}_0, e_1, \hat{v}_1, \ldots, e_\ell, \hat{v}_\ell)$, with $\hat{v} = \hat{v}_i$. The Extract-Maximal-Unitigs algorithm starts two walks $w_b$ and $w_f$ from $\hat{v}_i$, respectively through its $\mathtt{back}$ and $\mathtt{front}$, using the algorithm Walk-Maximal-Unitig. WLOG, assume that $e_i$ and $e_{i+1}$ are incident to the $\mathtt{front}$ and to the $\mathtt{back}$ of $\hat{v}_i$, respectively. Also, let $p_f = (\hat{v}_0, e_1, \ldots, e_i, \hat{v}_i)$, and $p_b = (v_i, e_{i+1}, \ldots, e_\ell, \hat{v}_\ell)$. First consider the case that $|p_b| > 1$, so $i < \ell$. Since the $\mathtt{back}$ of $\hat{v}_i$ is internal to $p$, it only has the edge $e_{i+1}$, encoded in the automaton $M_v$'s state. So $w_b$ must exit $\hat{v}_i$ using $e_{i+1}$, entering $\hat{v}_{i+1}$. Now each $\hat{v}_j$ $(i < j < \ell)$ being an internal vertex to $p$, it only has the unique edges $e_j$ and $e_{j+1}$, one per each side. So $w_b$ enters each $\hat{v}_j$ with $e_j$ and exits it with $e_{j+1}$, thus continuing on. And it is not possible for $w_b$ to deviate off $p$ without reaching $\hat{v}_\ell$, where it terminates finding $\hat{v}_\ell$ to be flanking. Besides, early termination at some internal $\hat{v}_j$ $(i < j < \ell)$ does not occur either, as Lemma 3 implies that no internal vertex is flanking. Thus $w_b$ traverses $p_b$ in its entirety. For the case when $|p_b| = 1$, $w_b$ terminates immediately finding $\hat{v}_i$ to be flanking. Thus in either case, $w_b$ extracts $p_b$ correctly.

By symmetry, $w_f$ extracts $p_f$ correctly. Therefore $p$ is correctly constructed by joining $p_f$ and $p_b$ at $\hat{v}_i$.

Since each $v \in \mathcal{V}$ is processed in this manner to compute its containing maximal unitig, CUTTLEFISH 2 correctly extracts the entire set of maximal unitigs of $G(\mathcal{R}, k)$. ■
